# Engineering transposon-associated TnpB-ωRNA system for efficient gene editing and disease treatment in mouse

**DOI:** 10.1101/2023.06.29.547137

**Authors:** Zhifang Li, Ruochen Guo, Xiaozhi Sun, Guoling Li, Yuanhua Liu, Xiaona Huo, Rongrong Yang, Zhuang Shao, Hainan Zhang, Weihong Zhang, Xiaoyin Zhang, Shuangyu Ma, Yinan Yao, Xinyu Liu, Hui Yang, Chunyi Hu, Yingsi Zhou, Chunlong Xu

**Affiliations:** Lingang Laboratory, Shanghai, China.; School of Life Sciences and Technology, ShanghaiTech University, Shanghai, China; Shanghai Center for Brain Science and Brain-Inspired Technology, Shanghai, China.; HuidaGene Therapeutics Inc., Shanghai, China.; Department of Biological Sciences, National University of Singapore, Singapore; Institute of Neuroscience, State Key Laboratory of Neuroscience, Key Laboratory of Primate Neurobiology, CAS Center for Excellence in Brain Science and Intelligence Technology, Shanghai Institutes for Biological Sciences, Chinese Academy of Sciences, Shanghai, China.; Department of Histoembryology, Genetics and Developmental Biology, Shanghai Key Laboratory of Reproductive Medicine, Shanghai JiaoTong University School of Medicine, Shanghai, China.

## Abstract

Transposon-associated ribonucleoprotein TnpB is known to be the ancestry endonuclease of diverse Cas12 effector proteins from type-V CRISPR system. Given its small size (409 aa), it is of interest to examine whether engineered TnpB could be used for efficient mammalian genome editing. Here, we showed that the gene editing activity of native TnpB in mouse embryos was already higher than previously identified small-sized Cas12f1. Further stepwise engineering of noncoding RNA (ωRNA or reRNA) component of TnpB significantly elevated the nuclease activity of TnpB. Notably, an optimized TnpB-ωRNA system could be efficiently delivered *in vivo* with single adeno-associated virus (AAV) and prevented the disease phenotype in a tyrosinaemia mouse model. Thus, the engineered miniature TnpB system represents a new addition to the current genome editing toolbox, with the unique feature of the smallest effector size that facilitate efficient AAV delivery for editing of cells and tissues.

## Introduction

The TnpB proteins represent a family of transposon-associated RNA-guided endonucleases. Recent biochemical studies^1, 2^ revealed that TnpB proteins are ancestry predecessors of Cas12 effector proteins in the type-V CRISPR system, and a 247-nucleotides (nt) noncoding RNA (termed ωRNA or reRNA) derived from the right end of transposon element is the required component for TnpB to recognize and cleave target DNA. The size of TnpB proteins, with ∼400 amino acid (aa) residues, is much smaller than their evolutionary progeny Cas12 proteins (mostly ∼1000 aa). Furthermore, in vitro studies^1, 2^ demonstrated that TnpB exhibited double-strand DNA cleavage activity guided by ωRNA. Therefore, there is potential for the use of this TnpB system in genome editing and therapeutic applications.

Gene editing using Cas9 or Cas12 systems has been widely used in animal models and recently applied in clinical trials. At present, AAV is the most commonly used delivery system and shown to be safe in gene therapy^3^. However, the maximal cargo size of AAV was limited to be 4.7 kilobase (kb) pairs, hindering efficient in vivo delivery of the large Cas9 or Cas12 protein via single AAV injection. This size problem is exacerbated in the use of base and prime editors comprising Cas9 (or Cas12) and fusion enzymes. Recent identification of compact CRISPR effector proteins Cas12f1 (∼500 aa)^4^ and Cas13 (∼700 aa)^5, 6^ represent potential solutions. However, the gene editing efficiency of Cas12f1 was relatively low^7–11^, whereas Cas13 exhibited collateral RNA cleavage activity with uncertain safety profile^12, 13^.

In the present study, we demonstrated that genome editing activity of TnpB was markedly higher than that of Cas12f1 in cultured cells and mouse embryos. To further optimize the TnpB system, we engineered TnpB-associated ωRNA in a stepwise manner to identify the optimal ωRNA variant with the shortest sequence length and elevated gene editing activity. Importantly, we showed that the optimized TnpB-ωRNA system could be effectively delivered *in vivo* via a single AAV injection in tyrosinaemia model mice, leading to the prevention of disease phenotype. Thus, we have shown the applicability of the engineered hypercompact TnpB for genome editing *in vivo*.

## Results

### TnpB exhibited gene editing activity higher than Cas12f1

Previous study has shown the endonuclease activity of several Cas12f1 orthologs from type V-U CRISPR family that have small sizes. As the ancestry enzyme of Cas12 proteins, TnpB (∼400 aa) represents the smallest programmable nuclease among common single effector Cas proteins, including SpCas9, LbCas12a, Un1Cas12f1, and IscB (**Fig. 1a**). However, the mammalian genome editing potential of TnpB remained to be fully characterized. Thus, we selected several genomic loci to evaluate the editing activity of TnpB (from *Deinococcus radiodurans,* ISDra2) in mouse embryos. First, we in vitro transcribed ωRNA that targets the mouse *Tyr* gene (**Fig. 1b**), and inject ωRNA together with TnpB mRNA into mouse embryos. The injected embryos were then transferred into surrogate female mice to generate gene-modified offspring. Since *Tyr* gene encodes the black coat color of C57/B6 mice, we estimated the efficiency of TnpB-induced gene disrupton by directly examining the coat color change in TnpB-injected mice. We found that TnpB treatment completely converted black coat color into albino white in all newborn mice (**Fig. 1c**). In contrast, similar embryo injection of Un1Cas12f1 together with sgRNA targeting the *Tyr* gene did not change the black coat color in the newborn mice (**Fig. 1c**), suggesting a much lower *Tyr* gene disruption efficiency of Un1Cas12f1 than that of TnpB. Further deep-sequencing for *Tyr* gene showed that 20% and 90% of indel mutations were induced by Un1Cas12f1 and TnpB, respectively (**Fig. 1b**). Although Cas12f1 and TnpB have different requirements for target adjacent motif (TAM, also known as PAM) that recognizes the target sequence, we have chosen the targeted sequence in *Tyr* gene to have 17-bp overlap (among 20 bp) for both enzymes (**Fig. 1b**). Thus, the higher editing efficiency of TnpB as compared to Cas12f1 was largely due to its endonuclease activity.

**Fig. 1.**
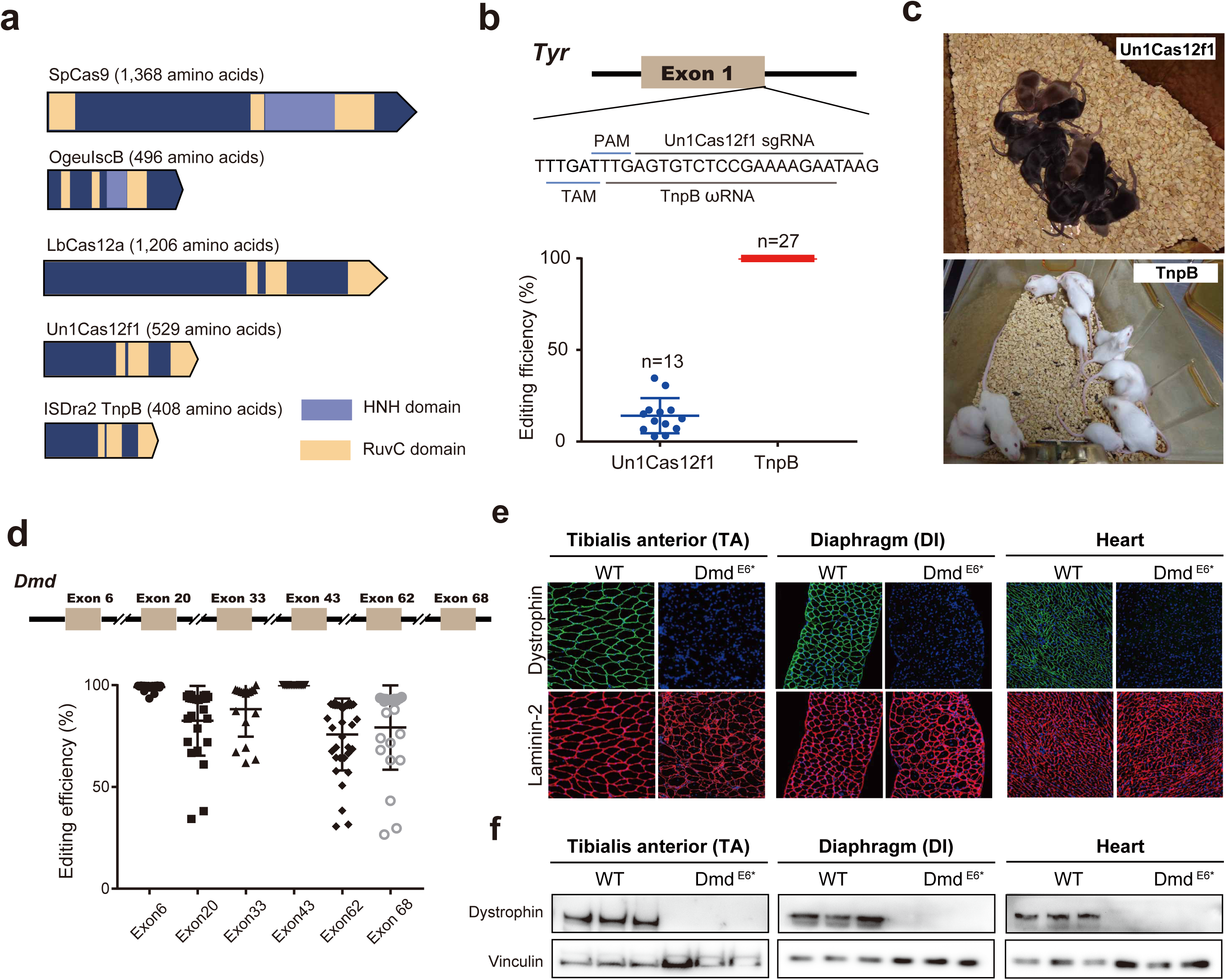
Mouse embryo injection of TnpB-ωRNA induced efficient gene editing. **a.** Characteristics of SpCas9, IscB, LbCas12a, Un1Cas12f1 and TnpB nucleases. **b.** Comparison of editing efficiency between TnpB and Cas12f1 on Tyr gene for gene modified mice. c. Coat color phenotype of *Tyr* gene modified mice by Un1Cas12f1 and TnpB. **d.** TnpB-mediated gene editing efficiency for *Dmd* gene. **e.** Dystrophin and laminin-2 immunostaining for TA, DI and heart muscle tissues in wildtype and *Dmd*-edited mice by TnpB. **f.** Western blotting of dystrophin and vinculin protein for three muscle tissues in wild-type and *Dmd*-edited mice by TnpB. Data are represented as means ± SEM. A dot represents a biological replicate. Significant differences between conditions are indicated by asterisk. Unpaired two-tailed Student’s t tests. * P < 0.05, *** P < 0.001, NS non-significant. Scale bars, 200 μm.

To further evaluate the gene editing activity of TnpB, we chose six additional loci in the mouse *Dmd* gene (**Fig. 1d, Supplementary Fig. 1**) for targeting in mouse embryos, by injecting ωRNA targeting these loci with TnpB mRNA. As shown by deep-sequencing results, TnpB exhibited an average of 90% editing efficiency for all six targeted loci in the *Dmd* gene (**Fig. 1d, Supplementary Fig. 1**). Furthermore, the gene editing outcome was verified by immunostaining of dystrophin protein encoded by *Dmd* gene that is specifically expressed in muscle tissues. In contrast to wildtype mice, TnpB-treated mice showed undetectable dystrophin expression in heart, Diaphragm (DI) and Tibialis anterior (TA) muscles (**Fig. 1e, Supplementary Fig. 2**), suggesting the complete disruption of *Dmd* gene by TnpB and ωRNA injection. Finally, these immunostaining results were confirmed by Western blotting of dystrophin protein of various muscle tissues (**Fig. 1f**). Consequently, rotarod and grip strength assessment of TnpB-treated DMD mice found functional dysfunction of muscle (**Supplementary Fig. 3**). Thus, our finding indicated more robust gene editing activity of TnpB than that of Un1Cas12f1 in mammalian tissues.

### Engineered TnpB-associated ωRNA with elevated editing efficiency

Cognate ωRNA scaffold associated with TnpB is 247 nt, much longer than sgRNA scaffold for most single effector Cas proteins. Previous findings reported that the sgRNA engineering could improve the performance of gene editing enzymes^14^. We thereby hypothesized that ωRNA truncation and optimization might be helpful for enhancing TnpB activity in mammalian cells. To this end, we predicted the secondary structure of ωRNA and formulated a stepwise strategy to truncate ωRNA (**Fig. 2a**). Based on the stem loops in predicted structure, we divided ωRNA into six segments, named as S1 to S6 for the truncation experiment (**Fig. 2b**). To facilitate screen of ωRNA variants, we designed a gene editing reporter with TnpB target DNA placed within a split and frameshifted GFP gene which could only be repaired after disruption of TnpB target sequence to express GFP (**Fig. 2a**). We tested the reporter with cognate ωRNA to prove the conditional activation of GFP after treatment of TnpB guided by ωRNA targeting frameshift mutation in GFP gene (**Fig. 2a**). At first, we deleted S1 to S6 one by one and run the reporter assay. It showed that only deletion of S4 and S6 ablated the activity of TnpB (**Fig. 2c**), suggesting the dispensable role of S1, S2, S3 and S5 for normal ωRNA function. Furthermore, sequence deletion of S1 slightly increase TnpB activity (**Fig. 2c**).

**Fig. 2.**
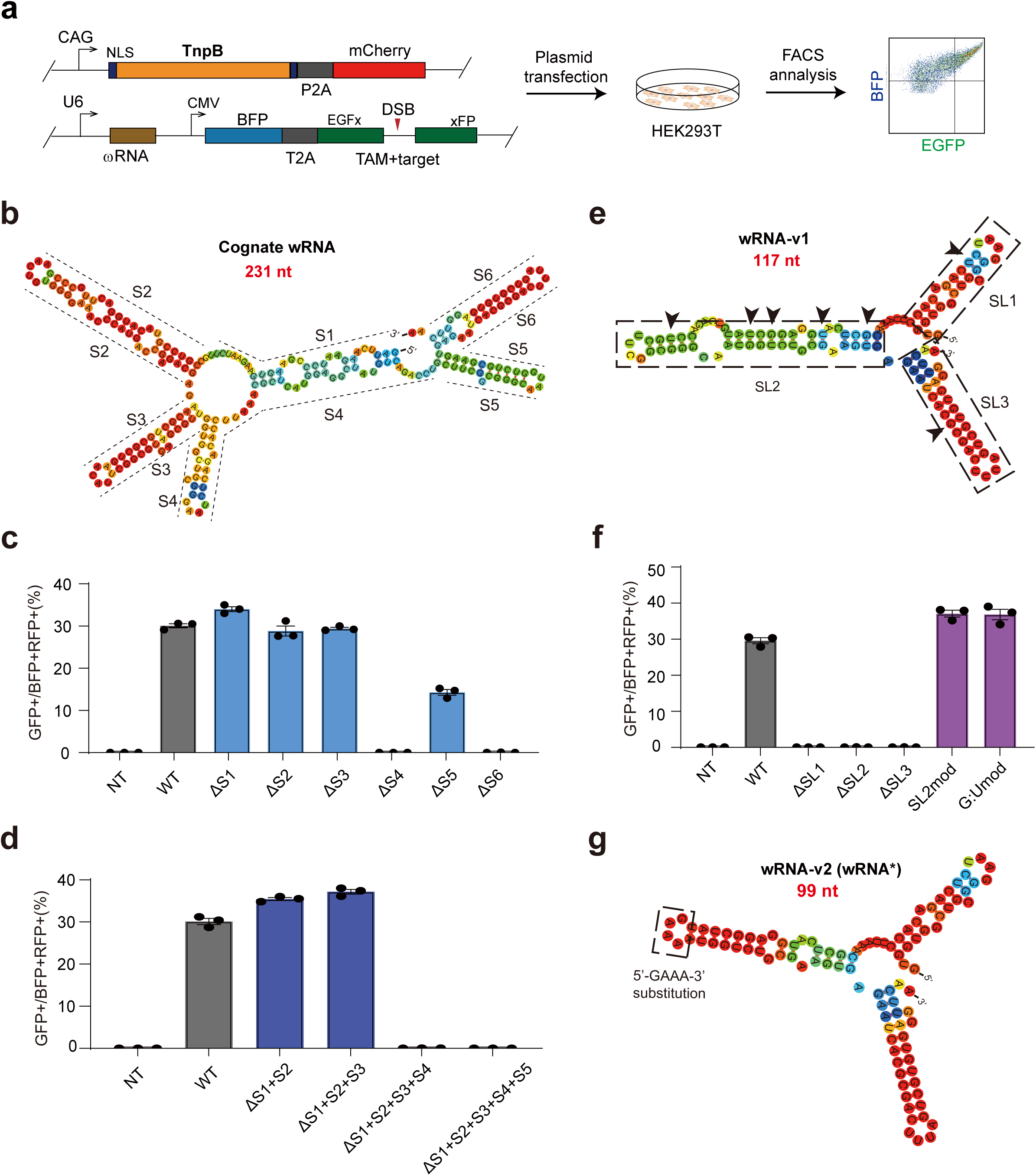
Stepwise engineering of TnpB-associated ωRNA improved gene editing efficiency. **a.** Reporter assay schematics of detecting cleavage activity of TnpB-ωRNA. **b.** Predicted secondary structure of cognate ωRNA (245 nt). Cognate ωRNA was divided into 6 segments, named from S1 to S6. **c.** Reporter assay results using engineered ωRNA by one-by-one truncation of S1 to S6. **d.** Reporter assay results with engineered ωRNA by different combined truncations of S1 to S5. **e.** Predicted secondary structure of a ωRNA variant with simultaneous truncation of S1, S2 and S3. **f.** Reporter assay results for ωRNA variants with different SL deletion and modifications. **g.** Predicted secondary structure of final optimized ωRNA variant. Data are represented as means ± SEM. A dot represents a biological replicate. Significant differences between conditions are indicated by asterisk. Unpaired two-tailed Student’s t tests. * P < 0.05, *** P < 0.001, NS non-significant.

To interrogate combined deletion effect of S1 to S6, we added S2 to S5 deletion in the S1 deletion variant of ωRNA to conduct reporter assay. It found that simultaneous deletion of S1, S2, and S3 in ωRNA-v1 not only supported the normal function of TnpB but also significantly increased the gene editing efficiency (**Fig. 2d**). These results implied that the ωRNA sequence from S4 to S6 dictated the enzymatic activity of TnpB. Secondary structure of ωRNA after combined truncation of S1, S2 and S3 showed typical stem loop conformations with three distinguishable and consecutive stem loop (SL) domains, termed as SL1, SL2 and SL3(**Fig. 2e**). To further determine the effect of these three SL domains on TnpB activity, we iteratively remove SL1, SL2 and SL3 for reporter test. In addition, we also generated two other ωRNA variants with partial deletion of SL2 subdomain or substitution of G:U with G:C pairs (**Fig. 2e**). We found that SL1, SL2 and SL3 are necessary for the normal function of TnpB since deletion variants lack of any single SL fully blocked the reporter activation (**Fig. 2f**). However, partial replacement of SL3 subdomain with 5’-GAAA-3’ loop sequence actually enhance the TnpB activity (**Fig. 2f**). G:C substitution for G:U pair exhibited no additive effect on the performance of TnpB (**Fig. 2f**). Based on these results, we finally identified an optimal ωRNA variant ωRNA-v2 or ωRNA* that improved TnpB performance. Predicted secondary structure of ωRNA* presented with three compact stem loop domains in contrast to loose organization of cognate ωRNA structure (**Fig. 2g**).

### Characterization of endogenous gene editing and off-target activity for TnpB-ωRNA system

To verify the reporter assay results for ωRNA*, we selected 14 endogenous genomic loci for further evaluation of gene editing performance in HEK293T (**Fig. 3a**). Among 14 human loci tested, 10 individual target sites showed significant increase of TnpB gene editing efficiency with ωRNA* compared with original ωRNA (**Fig. 3b**). Summary analysis of 14 loci also found significant improvement for TnpB using ωRNA* (**Fig. 3c**). To investigate broad improvement effect of ωRNA* in mammalian cells, we further performed the gene editing in mouse N2a cells targeting four disease relevant genes, including *Klkb1*, *Tyr*, *Hpd* and *Pcsk9*. It found that all genomic sites exhibited significantly increased gene editing efficiency for ωRNA* compared to original ωRNA (**Supplementary Fig. 4**). Quantitative analysis revealed two fold increase of gene editing efficiency in N2a for ωRNA* versus original ωRNA (**Supplementary Fig. 4**). In particular, ωRNA* even supported TnpB editing of some loci that are barely edited using cognate ωRNA (**Supplementary Fig. 4**). Therefore, we demosntrated the enhanced TnpB activity in mammalian cells via the identification of ωRNA* after stepwise engineering.

**Fig. 3.**
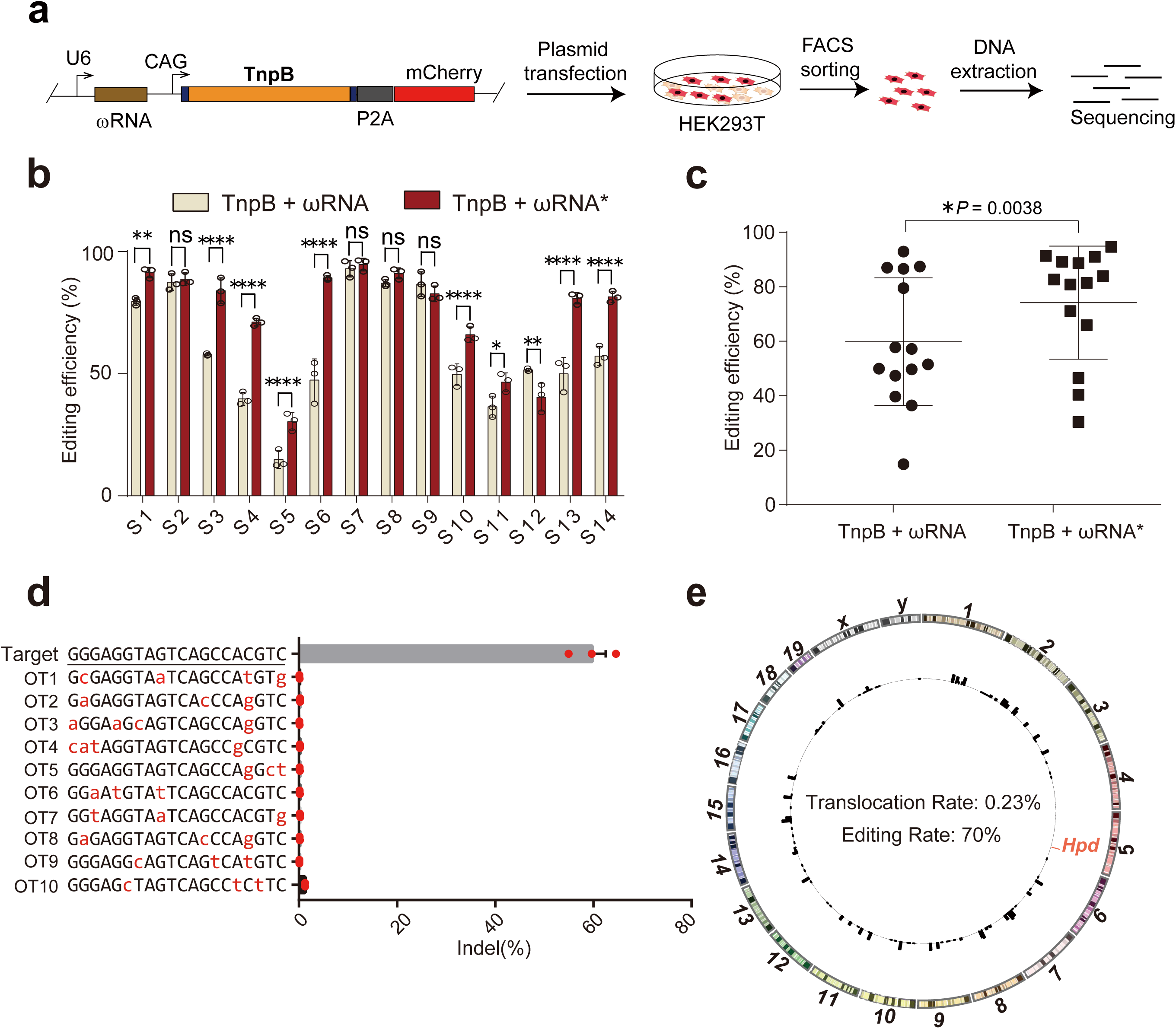
Characterization of endogenous gene editing activity and off-target effect with optimized TnpB-ωRNA system. **a.** The experimental workflow for detecting editing efficiency of original and optimized TnpB-ωRNA in HEK293T cells. **b.** Efficiency comparison results for 14 endogenous gene edited with original and optimized TnpB-ωRNA. **c.** Summary results for 14 endogenous genes editing efficiency. **d.** Off-target analysis for top predicted off-target genomic loci via Cas-OFFinder. **e.** Genome-wide off-target analysis with PEM-seq. Data are represented as means ± SEM. A dot represents a biological replicate. Significant differences between conditions are indicated by asterisk. Unpaired two-tailed Student’s t tests. * P < 0.05, *** P < 0.001, NS non-significant.

To examine the off-target effect of TnpB, we carried out prediction of potential off-target genomic loci with Cas-OFFinder^15^ for off-target analysis when designing ωRNA against a target site in Hpd gene (**Fig. 3d**). For the top 10 predicted off-target sites, no gene editing events was detected for Hpd-targeting TnpB-ωRNA (**Fig. 3d**). Furthermore, we also performed genome-wide off-target analysis by PEM-seq^16^ to identify potential tranlocation between on-target and off-target loci. Our PEM-seq results showed that there is no induction of translocation events related to gene editing of Hpd gene by TnpB-ωRNA treatment (**Fig. 3e**).

### Prevention of fatal liver disease with in vivo delivery of TnpB-ωRNA via single AAV

Given the hypercompact size of TnpB, it would greatly facilitate in vivo delivery via single AAV for gene editing therapy. To demonstrate the potential of TnpB in disease intervention, we chose the *Hpd* as therapeutic target for gene editing therapy of type I hereditary tyrosinaemia (HTI) in Fah^-/-^ mouse model. Adult Fah^-/-^ was administrated with AAV-TnpB or AAV-TnpB-ωRNA (**Fig. 4a**) and kept without NTBC drug, an HPD inhibitor. We observed that AAV-TnpB-ωRNA treated Fah^-/-^ mice was still alive after 75 days without NTBC but all untreated mice died at about 65 days(**Fig. 4b**). Furthermore, Fah^-/-^ mice subject to AAV-TnpB-ωRNA treatment gained body weight after experiencing a short period of weight loss (**Fig. 4c**). Contrarily, untreated mice exhibited rapid weight loss until death (**Fig. 4c**). Histological analysis found massive liver fibrosis in untreated mice whereas dramatically reduced fibrotic pathology in treated mice (**Fig. 4d**).

**Fig. 4.**
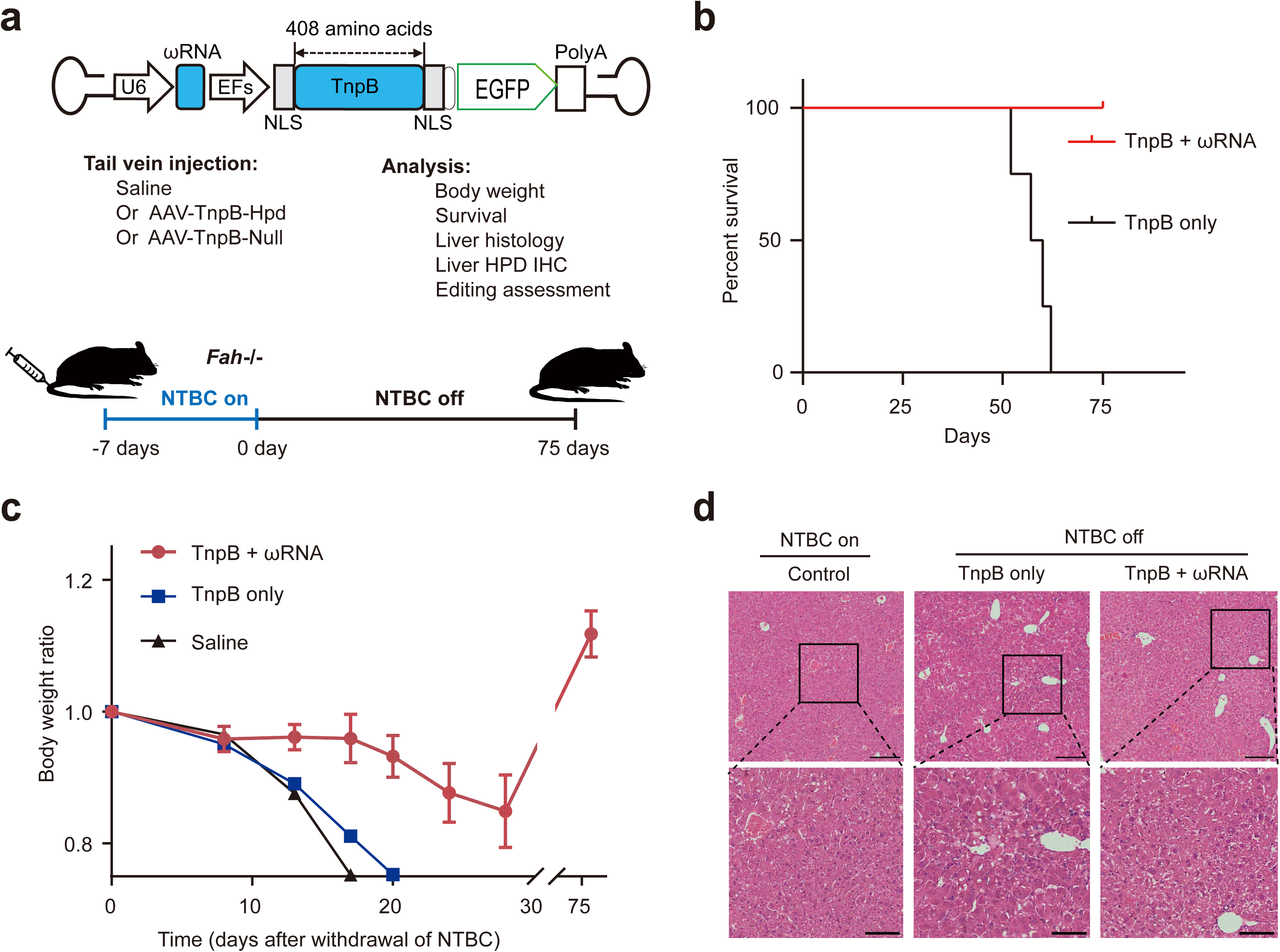
Prevention of fatal liver disease with in vivo delivery of TnpB-ωRNA via single AAV. **a.** Diagram of AAV-TnpB-ωRNA vector and gene therapy schematics in Fah-/-mouse model of type I hereditary tyrosinaemia. **b.** Survival curve for disease mice treated with AAV-TnpB-ωRNA or AAV-TnpB without ωRNA. **c.** Body weight change during the observation period for disease mice in different treatment groups. **d.** Histology analysis with H&E staining for mouse liver from different treatment groups. Data are represented as means ± SEM. A dot represents a biological replicate. Significant differences between conditions are indicated by asterisk. Unpaired two-tailed Student’s t tests. * P < 0.05, *** P < 0.001, NS non-significant. Scale bars, 200 μm.

Furthermore, we also analyzed the HPD expression in treated versus untreated mice. It showed the remarkable decrease of HPD positive liver region in AAV-TnpB-ωRNA treated mice (**Supplementary Fig. 5a, b**). To investigate the in vivo gene editing outcomes, we collected liver tissue from both treated and untreated mice. We only found 70% indel rate in AAV-TnpB-ωRNA treated mice but no editing events in non-treated animals (**Supplementary Fig. 5c, d**). Consistently, liver metabolic functions were significantly ameliorated after AAV-TnpB-ωRNA treatment as indicated by the blood biochemical profiling results of alanine aminotransferase (ALT), aspartate aminotransferase (AST), total bilirubin and tyrosine (**Supplementary Fig. 6**). Therefore, our results showed the proof-of-concept for applying TnpB in disease control via single AAV delivery in vivo.

## Discussion

Diverse CRISPR-Cas systems evolved from immune battle between microbe and mobile genetic elements (MGE), providing us abundant resources for the identification of gene editing enzymes^17^. In the past years, various single effector Cas proteins including Cas9^18^, Cas12^19^ and Cas13^20^ were found to deploy DNA or RNA editing activity in different organisms for both research and therapeutic purpose^21^. Recently, TnpB-like proteins, including IscB and TnpB associated with microbe transposon element, were identified to be active ancestry endonuclease for Cas9 and Cas12^1, 2^. Given the hypercompact size of TnpB and IscB, they are excellent candidates for developing miniature gene editing tools that would facilitate in vivo delivery via AAV. To this end, our present study demonstrated the potential of TnpB for robust genome editing in both cultured cells and animal tissues. Although Kim et al. recently reported engineering base editor from a 557-aa ‘TnpB’^22^, both Siksnys and Doudna group lately demonstrated that ‘TnpB’ used by Kim et al. study should be actually annotated as Cas12f1 that works as dimer unlike monomer TnpB^23, 24^. Thus, our work was the first study to extensively show the rational optimization of TnpB to achieve excellent in vitro and in vivo performance for gene editing. Furthermore, we also showed the effectiveness of TnpB based gene editing therapy to prevent fatal genetic disease in mouse model of tyrosineamia via in vivo single AAV delivery of TnpB and ωRNA. Interestingly, we performed stepwise truncation of cognate ωRNA to generate a ωRNA variant with short sequence and high efficiency. Our study represent a good start point to optimize TnpB or even IscB for more broad and convenient use in research and therapeutic scenario.

Endonuclease activity of TnpB was only shown with limited data in 2021 by Karvelis et al study^1^. Extensive characterization of TnpB activity in mammalian cell and tissue were currently needed. Our finding corroborated the results from Karvelis et al study, revealing unexpected higher activity for TnpB than Cas12f1 without further engineering. Moreover, we showed that deletion of 5’-end and partial internal sequence in ωRNA could enhance the gene editing performance of TnpB both in vitro and in vivo. Intriguingly, such deletion strategy was supported by two latest structural studies^25, 26^ of TnpB-ωRNA-DNA ternary complex published last month, suggesting the potential useful applicability of our ωRNA engineering strategy for more TnpB-like systems. In addition, the TnpB structure could accelerate the rational engineering of such compact enzyme with more demanding properties such as relaxed limitation of target-adjacent motif (TAM), enhanced editing activity and specificity etc.

Gene editing therapy was partly impeded by the limited AAV cargo capacity of only ∼4.7 kb considering the fact that common Cas9, Cas12 and their derived base or prime editors have protein size beyond 1000 aa^3, 27^. TnpB with less than 500 aa are highly desired gene editing enzymes for AAV delivery in vivo. Our results with TnpB in treating fatal tyrosineamia in mice signify the advantage of reducing gene editing cargo size despite the modest modification efficiency for Hpd target gene after TnpB-ωRNA optimization. Besides, compact TnpB size could permit using sophisticated regulatory sequences for switchable gene editing and reducing the AAV administration dose for high expression to enable safe therapeutic applications. Furthermore, our optimized ωRNA* variant with less than 100 nt would also be easy for synthesizing chemically modified ωRNA, which is very useful for ribonucleoprotein(RNP)-based gene editing applications.

Overall, our study demonstrated improved gene editing activity of TnpB via ωRNA engineering in cultured cells and showed its disease prevention ability in animal models, indicating the potential of hypercompact TnpB-ωRNA system as effective miniature gene editing modality for more AAV-based disease treatment in animal models and even human patients.

## Acknowledgements

We thank technical support from laboratory animal center (Y.D., J.S., T.Z.), optical imaging (L.T., K.S., W.L.) and gene-editing core (R.Y., X.H., X.Z.) facility in Shanghai Center for Brain Science and Brian-Inspired Technology as well as Lingang Laboratory. **Funding:** This work was funded by Lingang Laboratory (LG2023 to C.X.), and Shanghai City Committee of Science and Technology Project (22QA1412300 to C.X., 20ZR1466600 to X.H.). S.M. was funded by National Natural Science Foundation of China (32100641). H.Y. was funded by National Science and Technology Innovation 2030 Major Program (2021ZD0200900) (H.Y.), Chinese National Science and Technology major project R&D Program of China (2018YFC2000101) (H.Y.), Strategic Priority Research Program of Chinese Academy of Science (XDB32060000) (H.Y.), National Natural Science Foundation of China (31871502, 31901047, 31925016, 91957122 and 82021001) (H.Y.), Basic Frontier Scientific Research Program of Chinese Academy of Sciences From 0 to 1 original innovation project (ZDBS-LY-SM001) (H.Y.), Shanghai Municipal Science and Technology Major Project (2018SHZDZX05) (H.Y.), Shanghai City Committee of Science and Technology Project (18411953700, 18JC1410100, 19XD1424400 and 19YF1455100) (H.Y.) and the International Partnership Program of Chinese Academy of Sciences (153D31KYSB20170059) (H.Y.).

## Author contributions

Z.L., R.G. and C.X. jointly conceived the project and designed experiments. Y.Z. and C.X. supervised the whole project. Z.L. and G.L. generated mouse model. Z.L. and R.G. designed vectors, performing in vitro experiments and scanning confocal imaging. X.H. and X.S. assisted with construction plasmids and cell culture. R.Y. and X.Z. prepared AAV virus. R.G., Z.L., Z.S. and G.L. performed in vivo virus injection, tissue dissection, histological immunostaining and liver function experiments. Y.L. and Y.Z. performed bioinformatics analysis. R.G., G.L. and X.H. assisted with tissue dissection, immunostaining and animal breeding. Z.L., R.G., G.L., C.H., Y.Z. and C.X. analyzed the data and organized figures. Z.L., C.H., Y.Z. and C.X. wrote the manuscript with data contributed by all authors participated in project.

## Data and materials availability

Deep-seq data is deposited to the GEO repository under accession number PRJNA963402 and plasmids are available from the corresponding authors upon request.

## Supplementary Materials

Materials and Methods

Figures S1 to S6

Tables S1 to S3

Supplementary materials for

## Materials and Methods

### Study approval

The objectives of the present study were to show proof-of-concept for in vivo TnpB-mediated gene editing in wildtype and disease mice. All animal experiments were performed and approved by the Animal Care and Use Committee of Shanghai Center for Brain Science and Brian-Inspired Technology, Shanghai, China.

### Plasmid constructions

The pCBh-TnpB-hU6-BpiI plasmid encoded a human codon-optimized TnpB driven by CBh promoter, and hU6-driven ωRNAs with *Bpi*I cloning site. The sgRNA and ωRNA were designed suitable for Un1Cas12f1 and TnpB, then synthesized as DNA oligonucleotides and cloned into pCBh-Un1Cas12f1 or pCBh-TnpB to get the CRISPR targeting plasmids.

### Cell culture, transfection and flow cytometry analysis

HEK293T were maintained in Dulbecco’s modified eagle medium (DMEM) (Gibco, 11965092) supplemented with 10% fetal bovine serum at 37 °C and 5% CO_2_ in a humidified incubator. For sgRNA screening, CRISPR targeting plasmids and reporter were co-transfected using polyethylenimine (PEI) transfection reagent. After transfected cells were cultured with 48 hours, we carefully resuspended the cell pellet, and then analyzed or sorted by BD FACSAria II. Flow cytometry results were analyzed with FlowJo X (v.10.0.7).

### In vitro transcription of TnpB and ωRNA

TnpB mRNA was transcribed using the mMESSAGE mMACHINE T7 Ultra Kit (Invitrogen, AM1345). T7 promoter was added to ωRNA template by PCR amplification of pCX2280 using primer F and R. The PCR products purified with Omega gel extraction Kit (Omega, D2500-02), templates were transcribed using the MEGAshortscript Kit (Invitrogen, AM1354). The TnpB mRNA and ωRNA were purified by MEGAclear Kit (Invitrogen, AM1908), eluted with RNase-free water and stored at -80°C.

### Zygote injection and embryo transplantation

Eight-week-old B6D2F1 female mice were super ovulated and mate with B6D2F1 male mice, and fertilized embryos were collected from oviduct. The mixture of TnpB mRNA(50 ng/µL) and ωRNA (100 ng/µL) was injected into the cytoplasm of fertilized eggs using a FemtoJet microinjector(Eppendrof). The injected embryos were cultured in KOSM medium with amino acids at 37°C under 5% CO_2_ in a humidified incubator overnight and then transferred into oviducts of pseudo-pregnant ICR foster mothers at 0.5-d.p.c.

### AAV virus production

The adeno-associated virus 8 (AAV8) serotype was used in this study. The TnpB plasmids with ωRNA was sequenced before packaging into AAV8 vehicle, and the AAV vectors were packaged by transfection of HEK293T cell with helper plasmids. The virus titer was 5 × 10^13^ (AAV-TnpB), and 5 × 10^13^ (AAV-TnpB-ωRNA) genome copies/mL as determined by qPCR specific for the inverted terminal repeat.

### Gene editing treatment for tyrosinaemia mouse model

Mice were housed in a barrier facility with a 12-hour light/dark cycle and 18– 23 °C with 40–60% humidity. Diet and water were accessible at all times. Fah^-/-^ mice were kept on 10mg/L NTBC (Sigma-Aldrich, Cat. No. PHR1731) in drinking water when indicated. For hydrodynamic liver injection, AAV8 (4 × 10^11^ vg/mouse) in 200 μl saline were injected via the tail vein into 8-10 weeks old male and female mice. Mice were kept off NTBC water at 7 days post injection, and their body weights were recorded every 3-5 days. Mice were harvested at 75 days after NTBC water withdrawal for histology and DNA analysis. Control mice off NTBC water were harvested when reaching >20% weight loss.

### Histological analysis and Serum biochemistry

Liver tissues were harvested, and sections were fixed in 4% PFA overnight. The following antibodies were used: anti-HPD antibody (SantaCruz, sc-390279; dilution 1:100), anti-P21 antibody (Abcam, ab109199; dilution 1:200). Immunohistochemistry, immunofluorescence and hematoxylin and eosin (H&E) staining were performed by the standard procedures. Blood was collected using retro-orbital puncture before mice was sacrificed. ALT, AST, tyrosine and bilirubin levels in serum were determined using diagnostic ELISA Kits (Abcam, HWRK chem).

### Targeted deep sequencing

To analyze TnpB targeting efficiency, the DNA of successfully transfected cells or AAV8 treatment tissues were extracted with TIANamp Genomic DNA Kit(TIANGEN,) according to the manufacturer protocol. DNA was amplified with Phanta max super-fidelity DNA polymerase (Vazyme, P505-d1) for Sanger or deep sequencing methods. And deep sequencing libraries were used to add Illumina flow cell binding sequences and specific barcodes on the 5′ and 3’ end of the primer sequence. The products were pooled and sequenced with 150 paired-end reads on an Illumina Hiseq instrument. FASTQ format data were analyzed using the Cutadapt (v.2.8)41 according to assigned barcode sequences. CRISPResso2 was used for gene editing analysis^28^.

### PEM-seq analysis

Genome-wide off-target analysis was performed following PEM-seq protocol previously described^16^. The 20 μg genomic DNA from TnpB edited or control samples were fragmented with Covaris sonicator to generate 300-700 bp DNA. DNA fragments was tagged with biotin at 5’-end by one-round PCR extension using a biotinylated primer, primer leftover removed by AMPure XP beads and purified by streptavidin beads. The single-stranded DNA on streptavidin beads is ligated with a bridge adapter containing 14-bp random molecular barcode, and PCR product was generated via nested PCR to enrich DNA fragment containing the bait DSB events and tagged with illumine adapter sequences. The prepared sequencing library was sequenced by Hi-seq 2500 with 150 bp pair-end reads. PEM-seq data analysis was performed using PEM-Q pipeline with default parameters.

### Statistical analysis

The number of independent biological replicates were shown in the figure legend. The data are presented as means ± SEM. Differences were assessed using unpaired two-tailed Student’s t tests. Differences in means were considered statistically significant at *P* < 0.05.

## Supplementary figures and legend

**Fig. S1.**
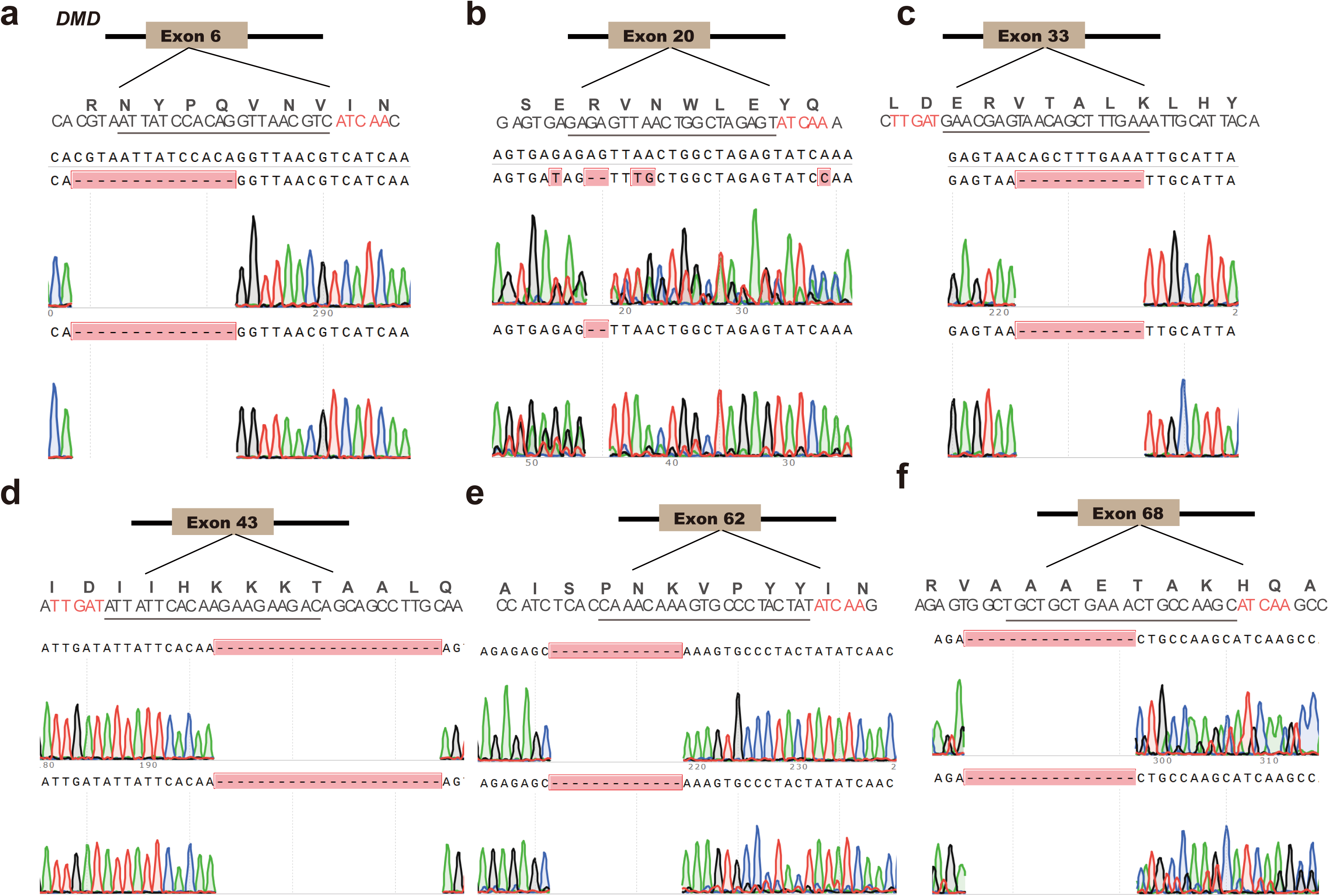
Transcriptional genotyping of Dmd-edited mice with RT-PCR. **a-f**. RT-PCR and sequencing results for muscle from individual mouse edited by TnpB in exon 6, 20, 33, 43, 62 and 68 of *Dmd* gene.

**Fig. S2.**
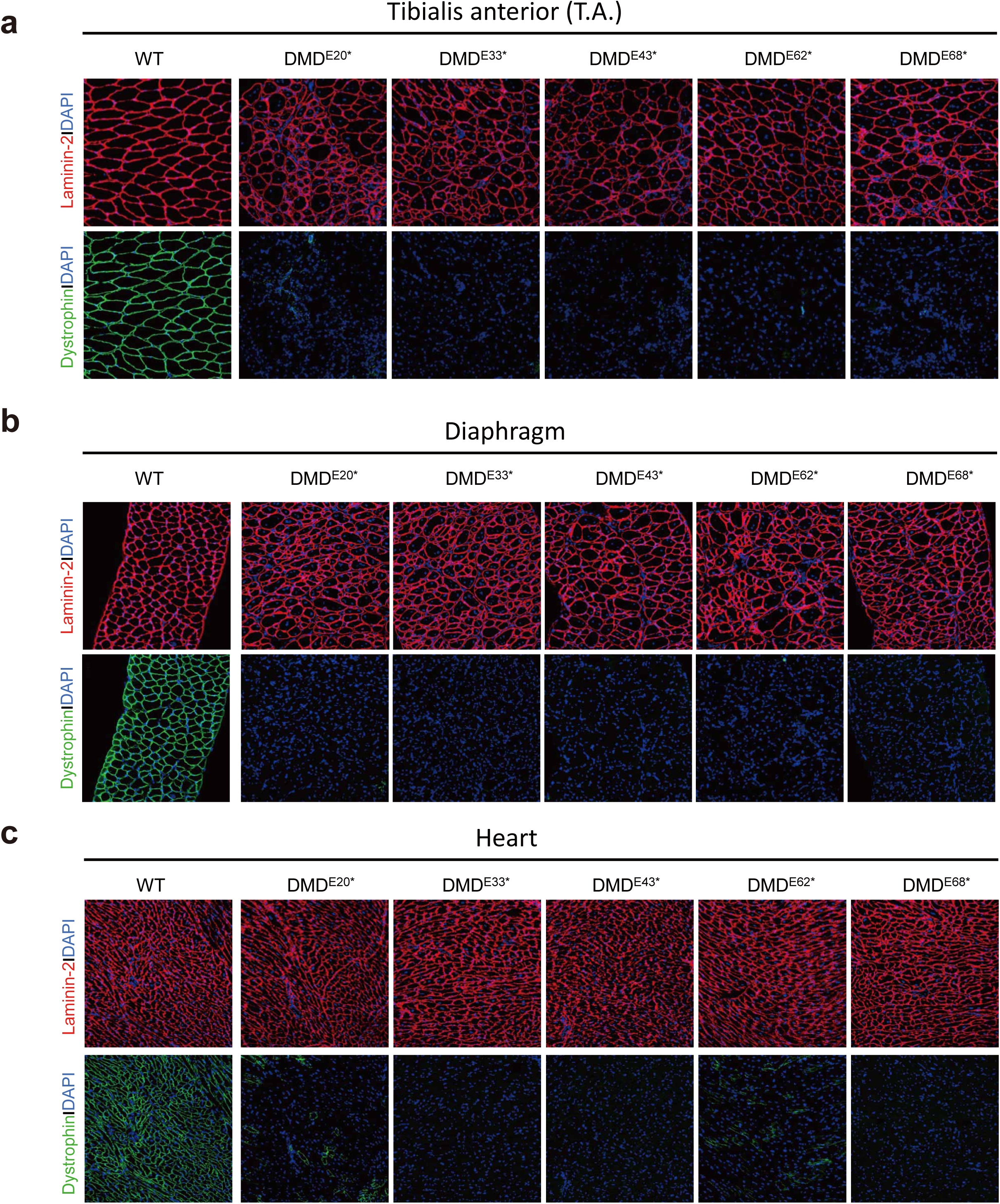
Dystrophin and laminin-2 immunostaining results for TA, DI and heart muscle in Dmd-edited mice. **a-c**. Immunostaining of dystrophin and laminin-2 in TA, DI and heart muscle from mice edited by TnpB in exon 6, 20, 33, 43, 62 and 68 of *Dmd* gene. Scale bars, 200 μm.

**Fig. S3.**
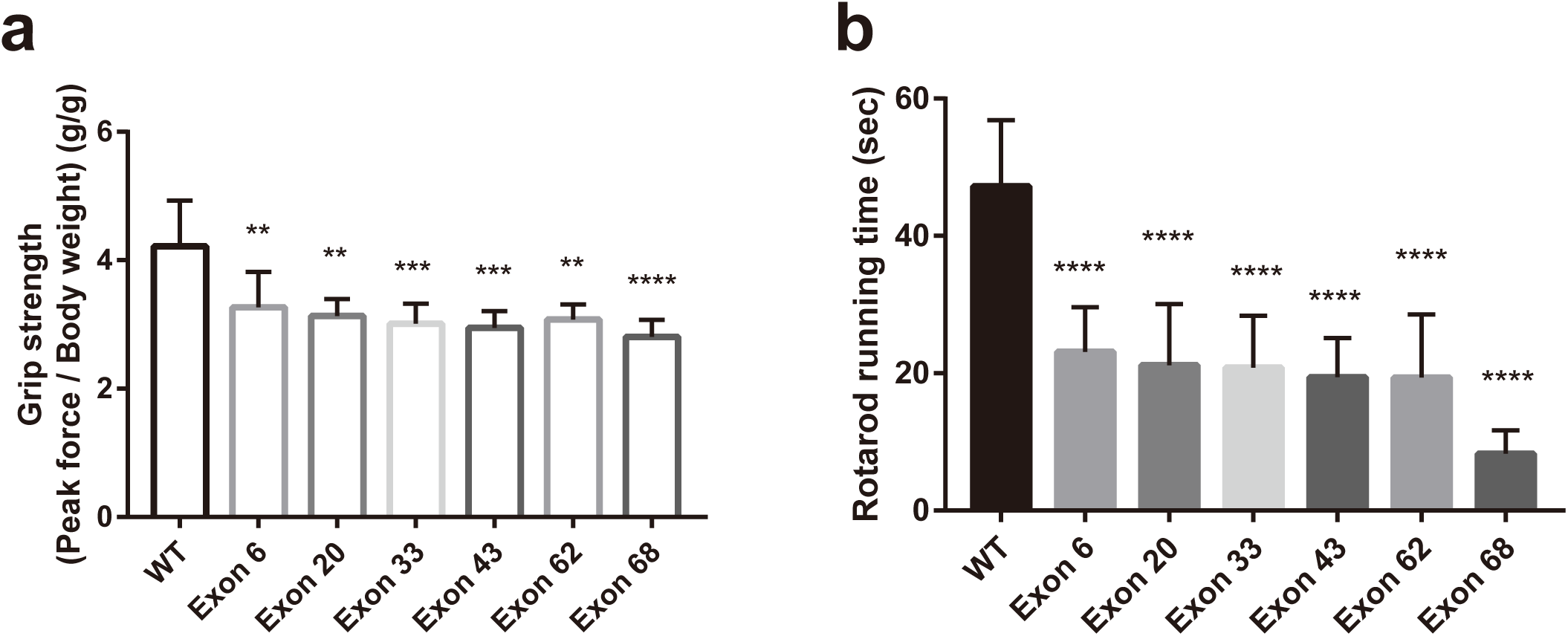
Grip strength and rotarod test for *Dmd*-edited mice. **a.** Forelimb grip strength analysis results for wildtype and *Dmd* mutant mice. **b.** Rotarod running time analysis results for wildtype and *Dmd* mutant mice. Data are represented as means ± SEM. A dot represents a biological replicate. Significant differences between conditions are indicated by asterisk. Unpaired two-tailed Student’s t tests. * P < 0.05, *** P < 0.001, NS non-significant.

**Fig. S4.**
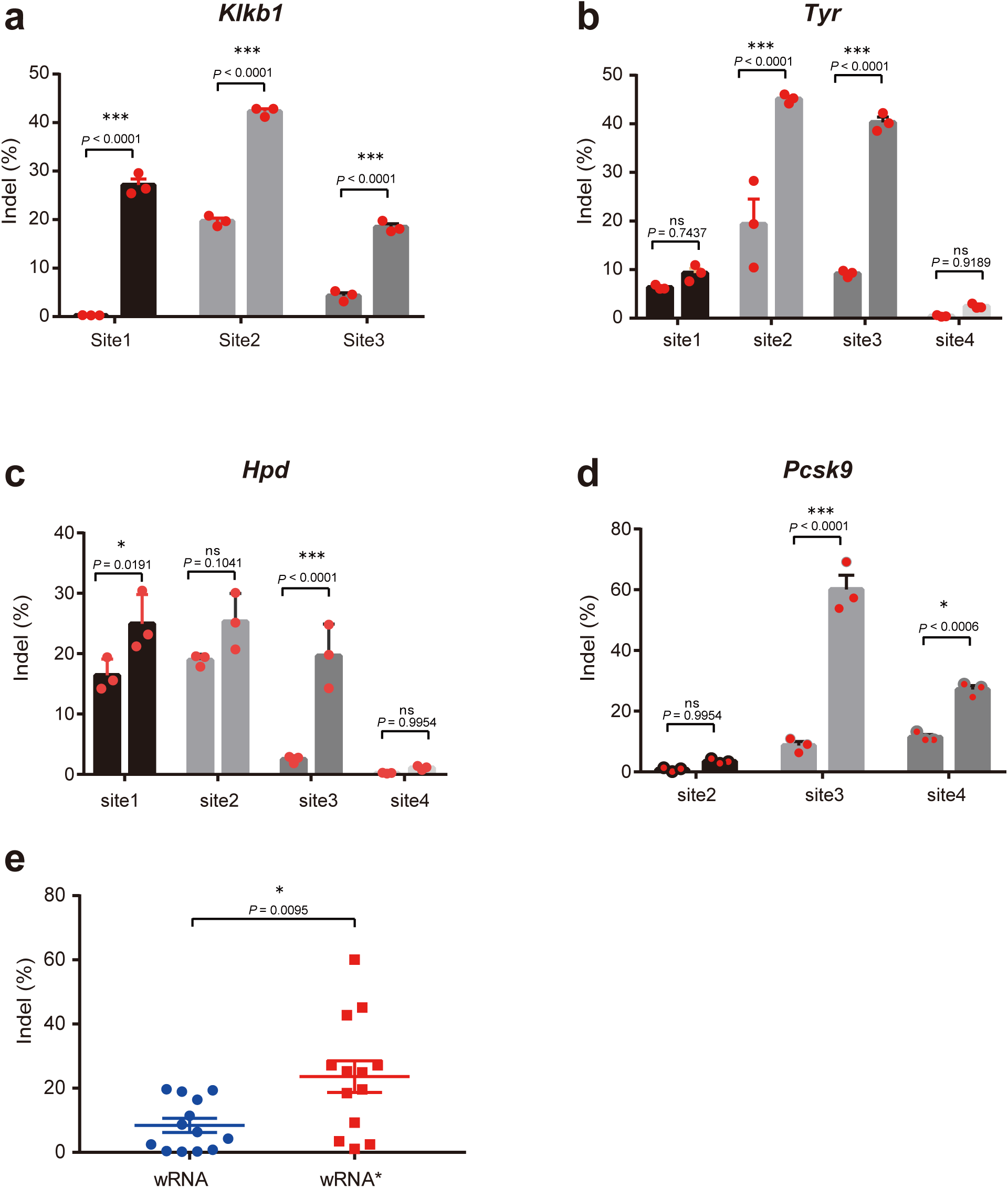
Characterization of gene editing activity for engineered TnpB-ωRNA system in mouse N2a cells. **a-d.** Efficiency comparison using cognate and engineered ωRNA for mouse *Klkb1, Tyr, Hpd*, and *Pcsk9* gene editing. **b.** Summary statistic results for gene editing activity characterization of cognate and engineered ωRNA in N2a. Data are represented as means ± SEM. A dot represents a biological replicate. Significant differences between conditions are indicated by asterisk. Unpaired two-tailed Student’s t tests. * P < 0.05, *** P < 0.001, NS non-significant.

**Fig. S5.**
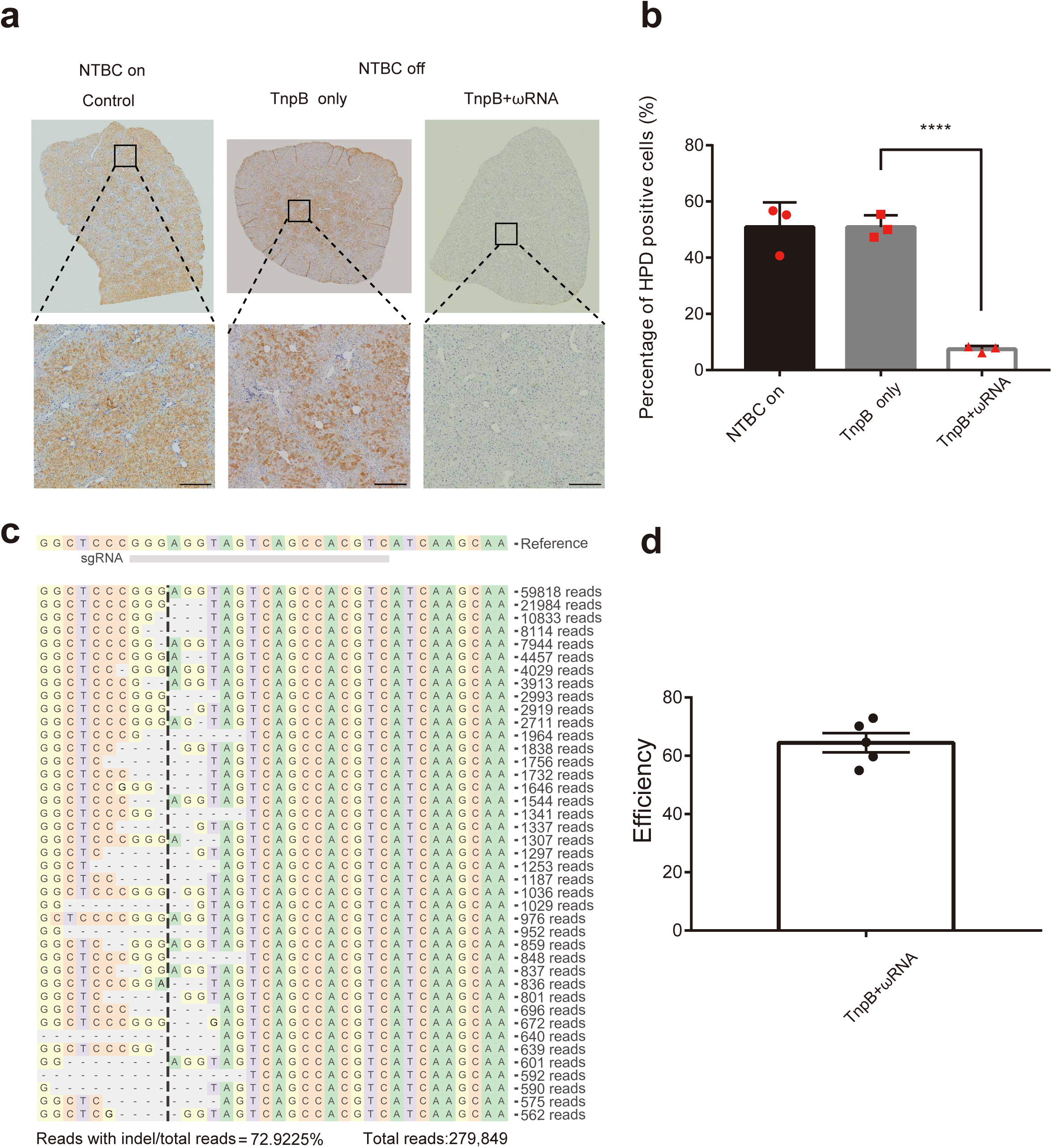
Gene editing and immunostaining analysis for HPD in AAV-TnpB-ωRNA treated mouse liver. **a.** Hpd immunostaining analysis in Fah^-/-^ mice treated with or without AAV-TnpB-ωRNA. **b.** Deep-seq results for *Hpd* gene editing by AAV-TnpB.

**Fig. S6.**
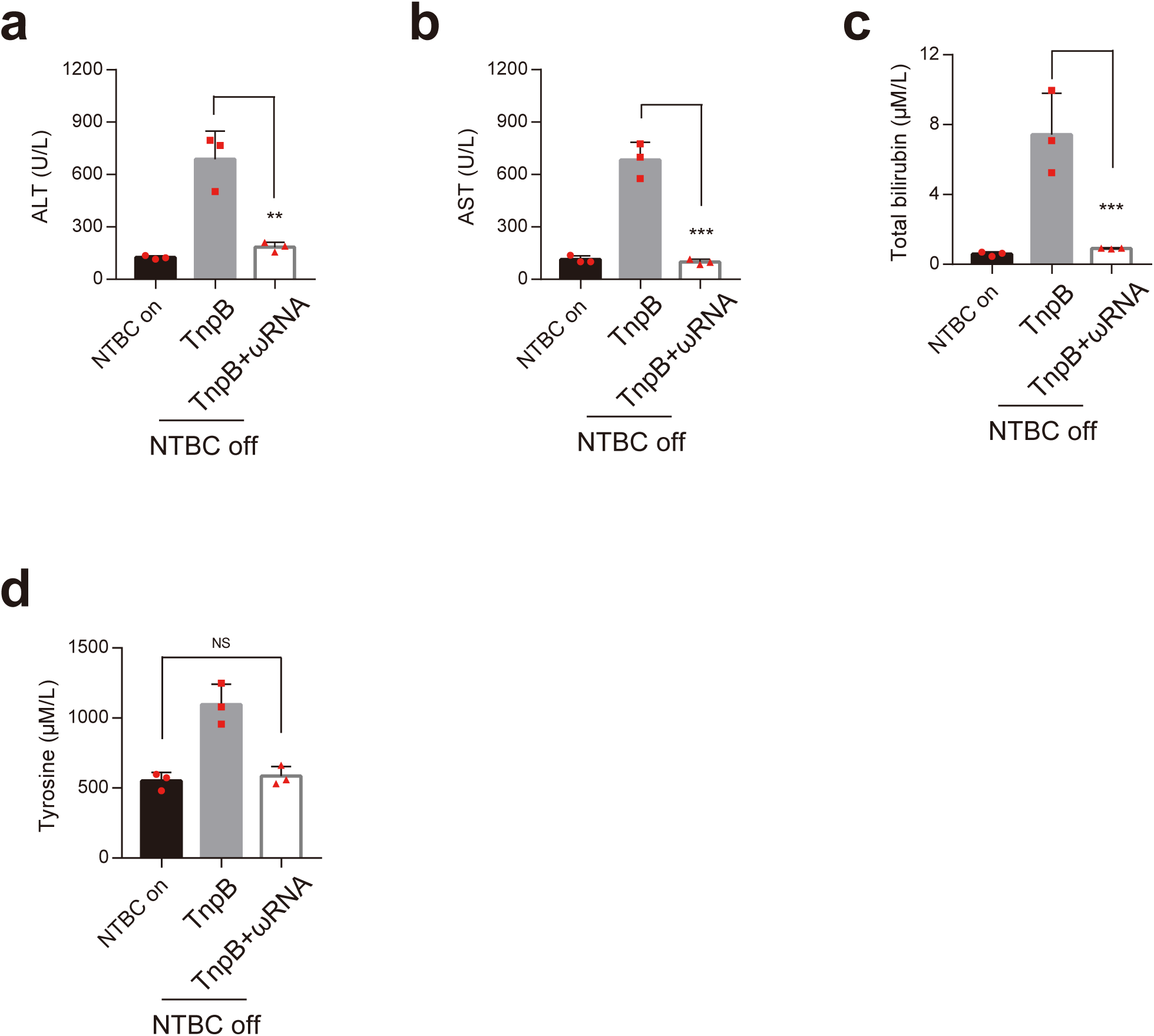
Serum biochemical analysis for AAV-TnpB-ωRNA treated mouse liver. **a-d.** Biochemical analysis of serum indicators for liver metabolic function in TnpB-treated or untreated mice (n=3). Liver damage markers alanine aminotransferase (ALT), aspartate aminotransferase (AST), total bilirubin and tyrosine were measured in peripheral blood from Fah^-/-^ mice injected with AAV-TnpB without or with ωRNA (NTBC off, day 30). Fah^-/-^ mice on NTBC water (NTBC on) served as a control. Data are represented as means ± SEM. A dot represents a biological replicate. Significant differences between conditions are indicated by asterisk. Unpaired two-tailed Student’s t tests. * P < 0.05, *** P < 0.001, NS non-significant.

**Supplementary Table S1.**
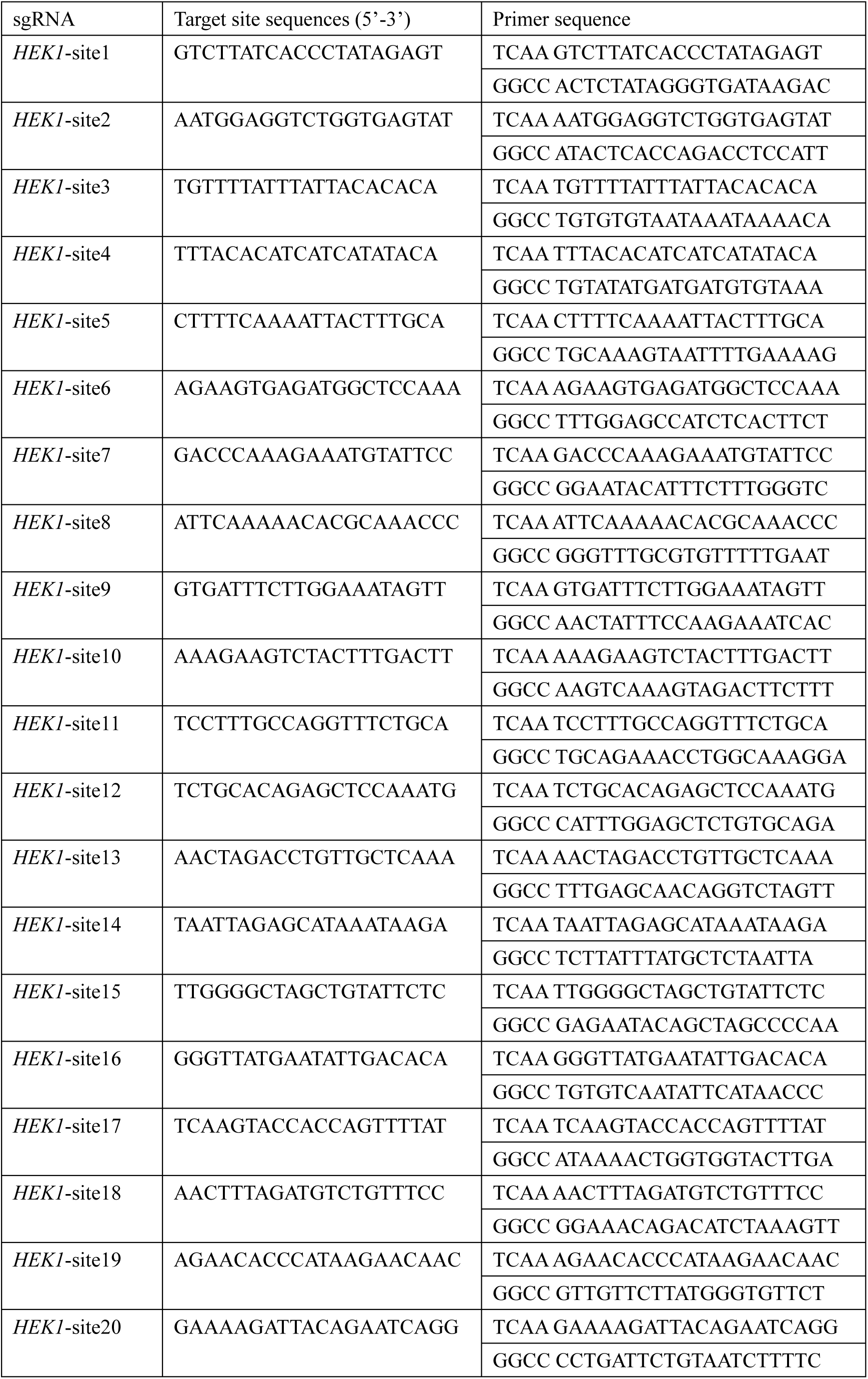
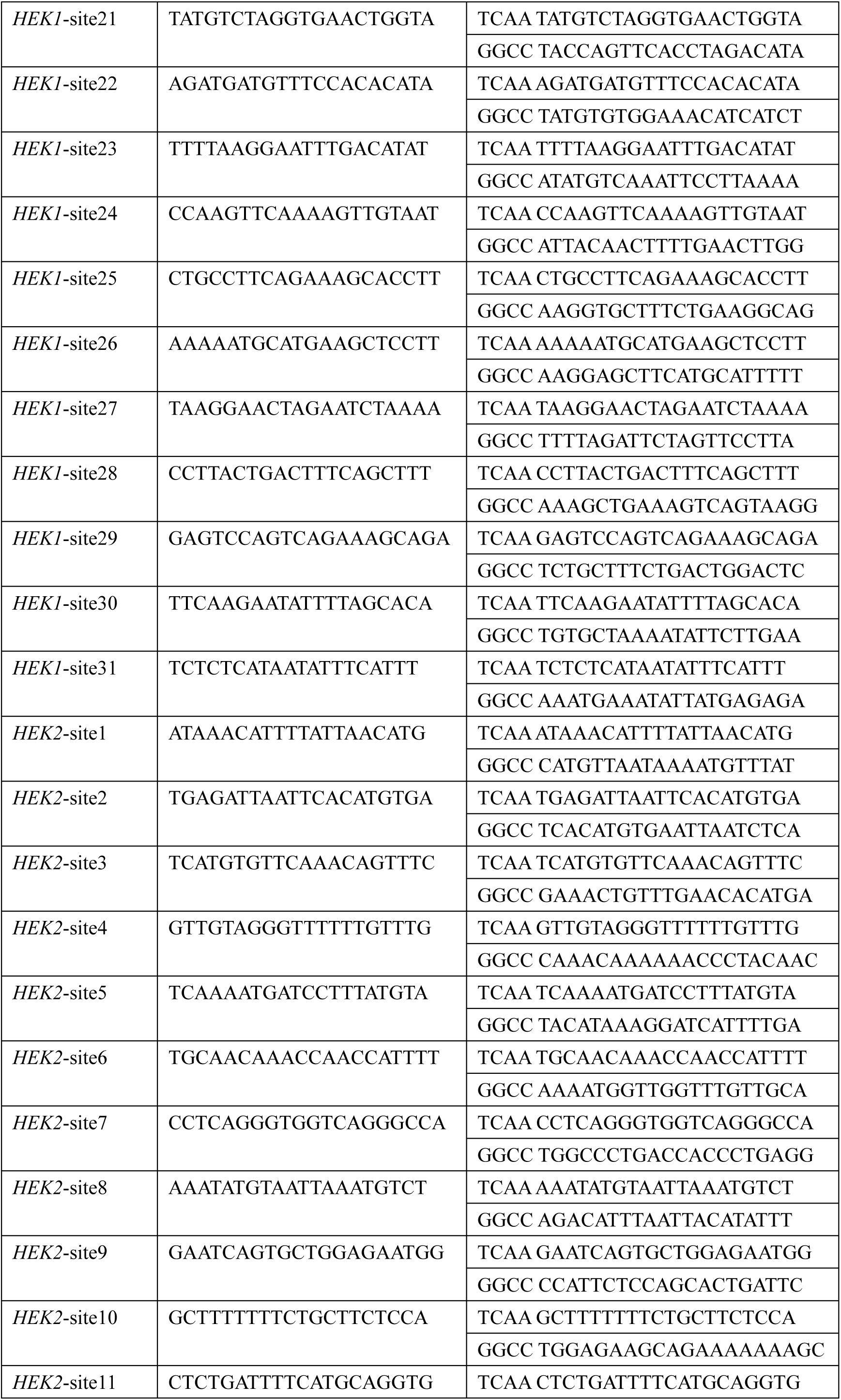
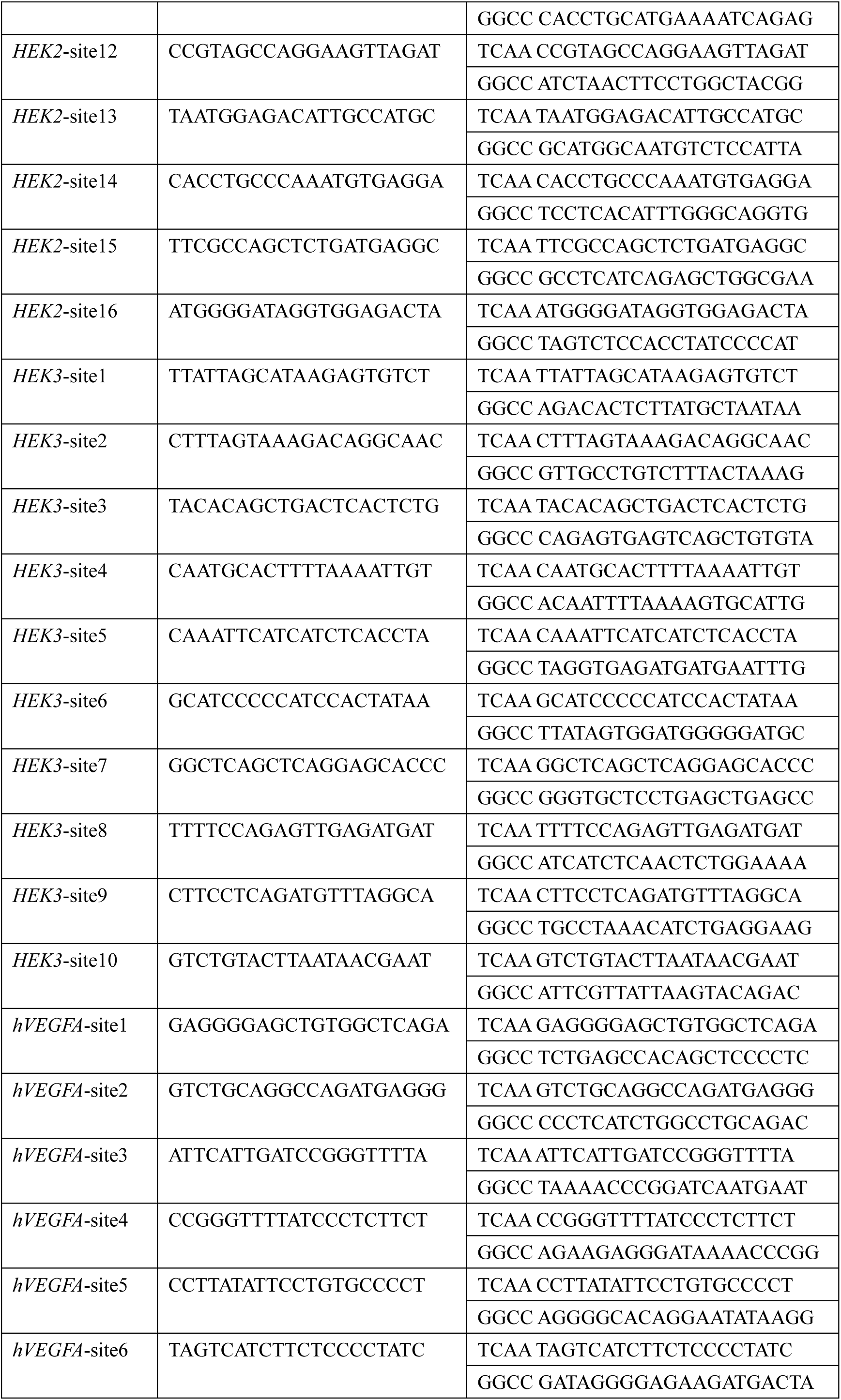
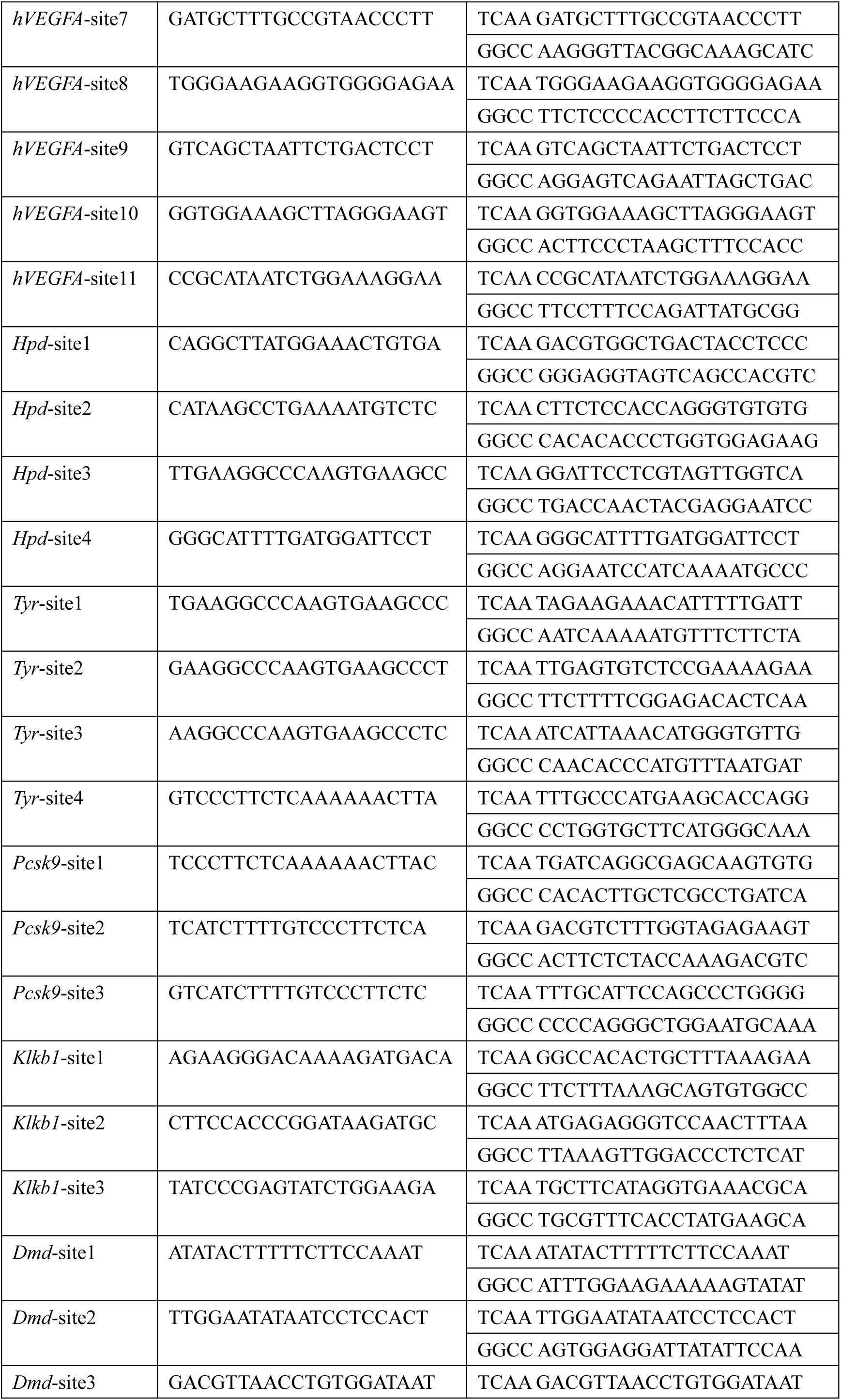
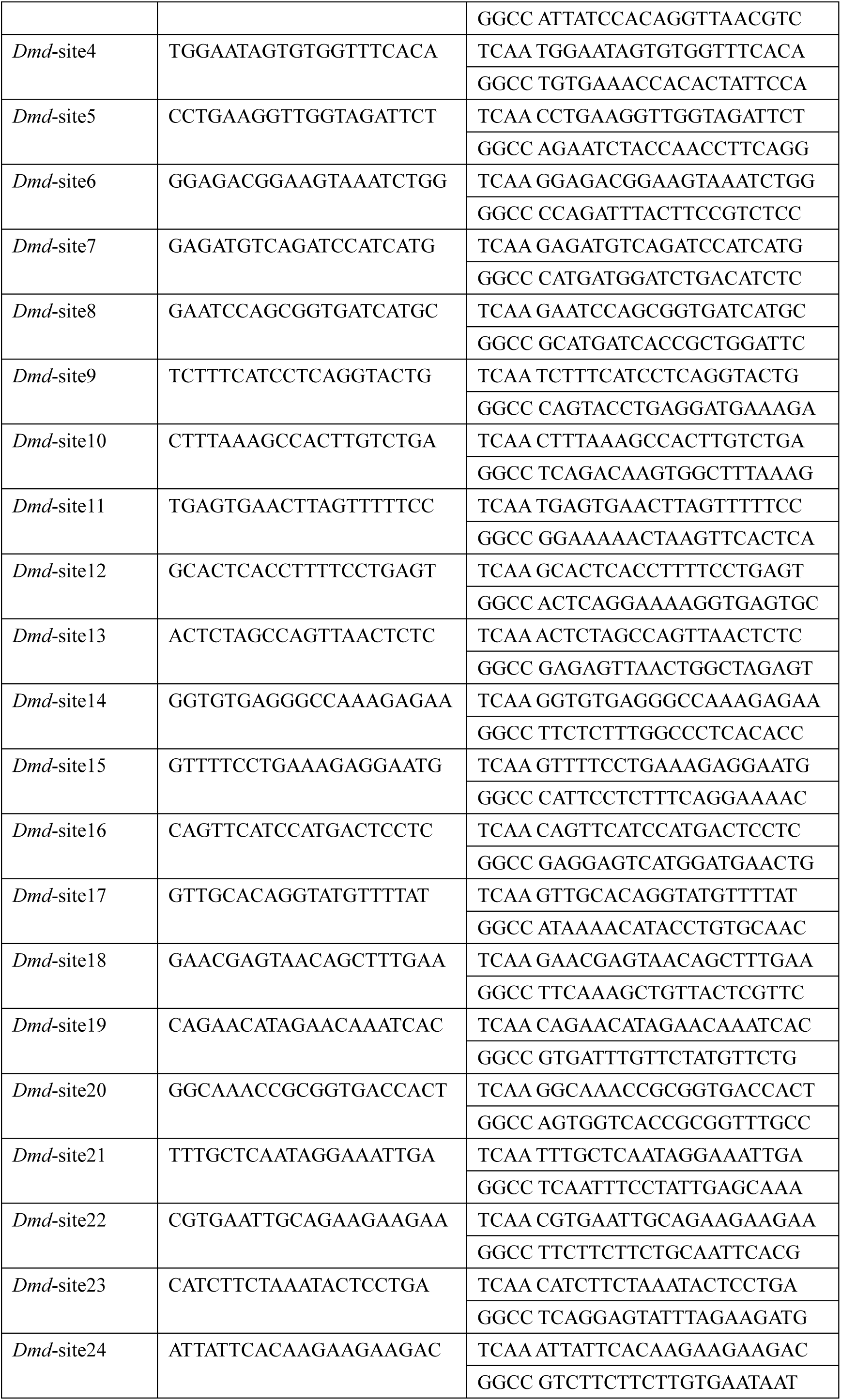
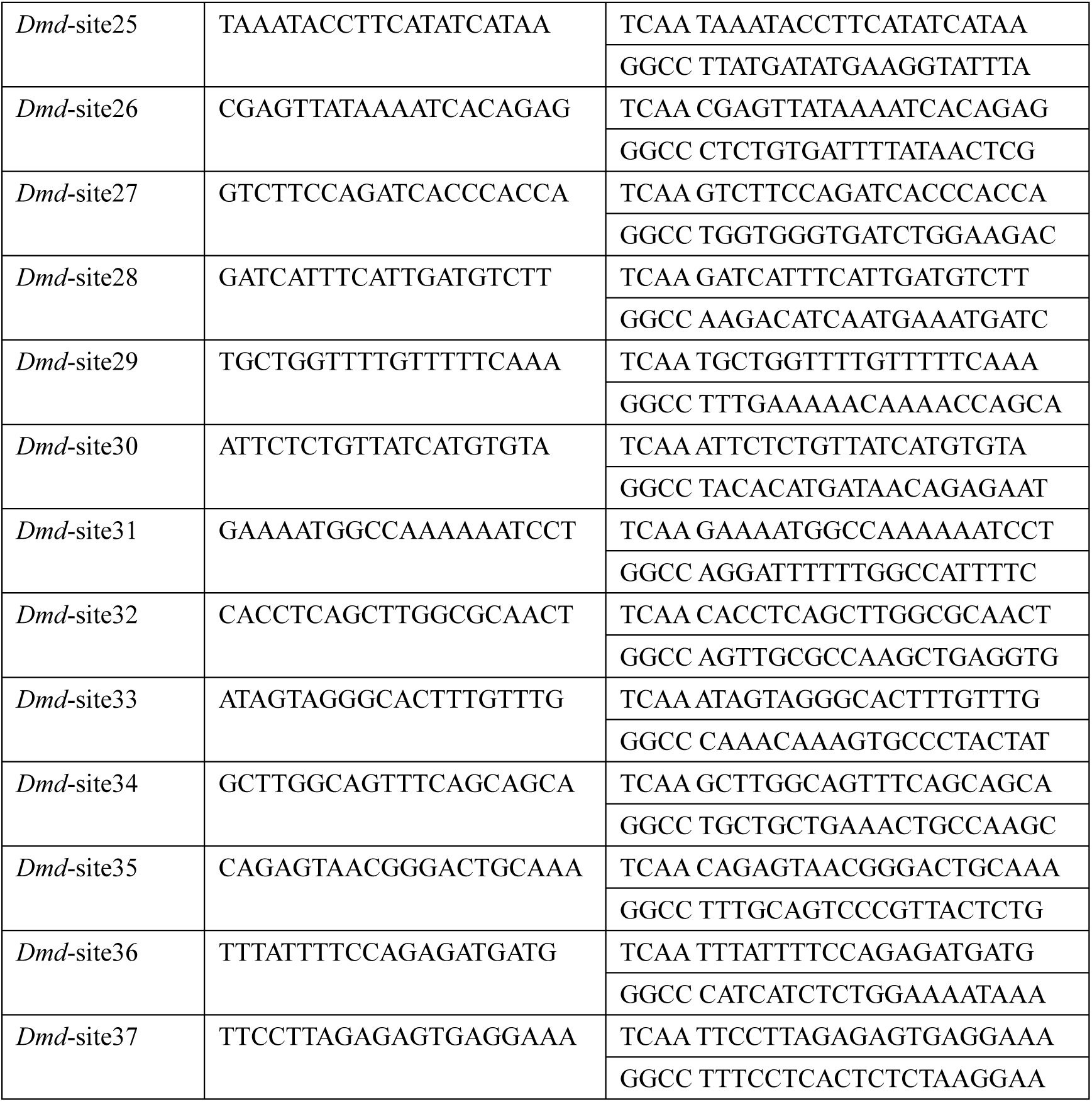
Target sgRNA and primer sequence.

**Supplementary Table S2.**
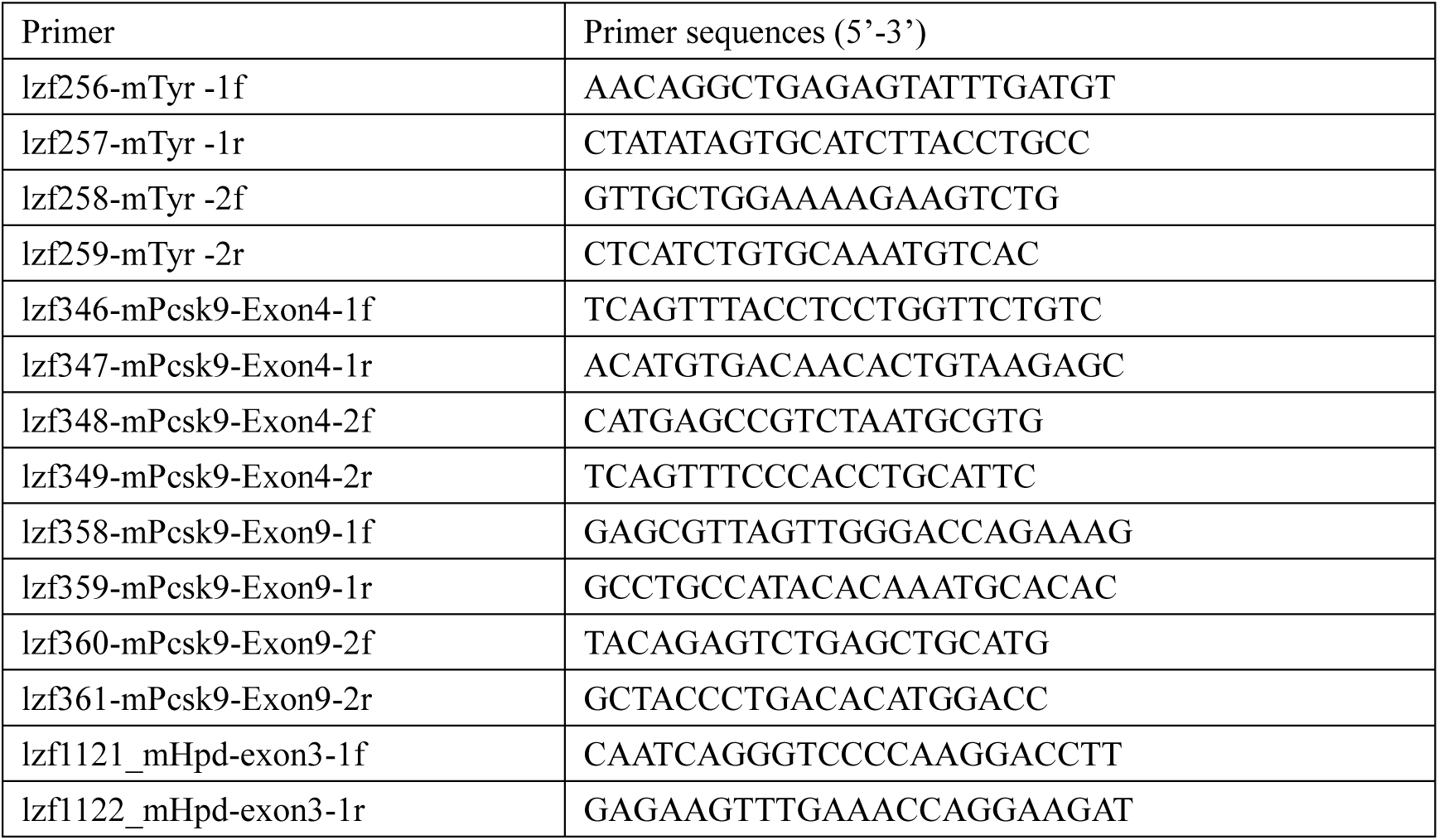
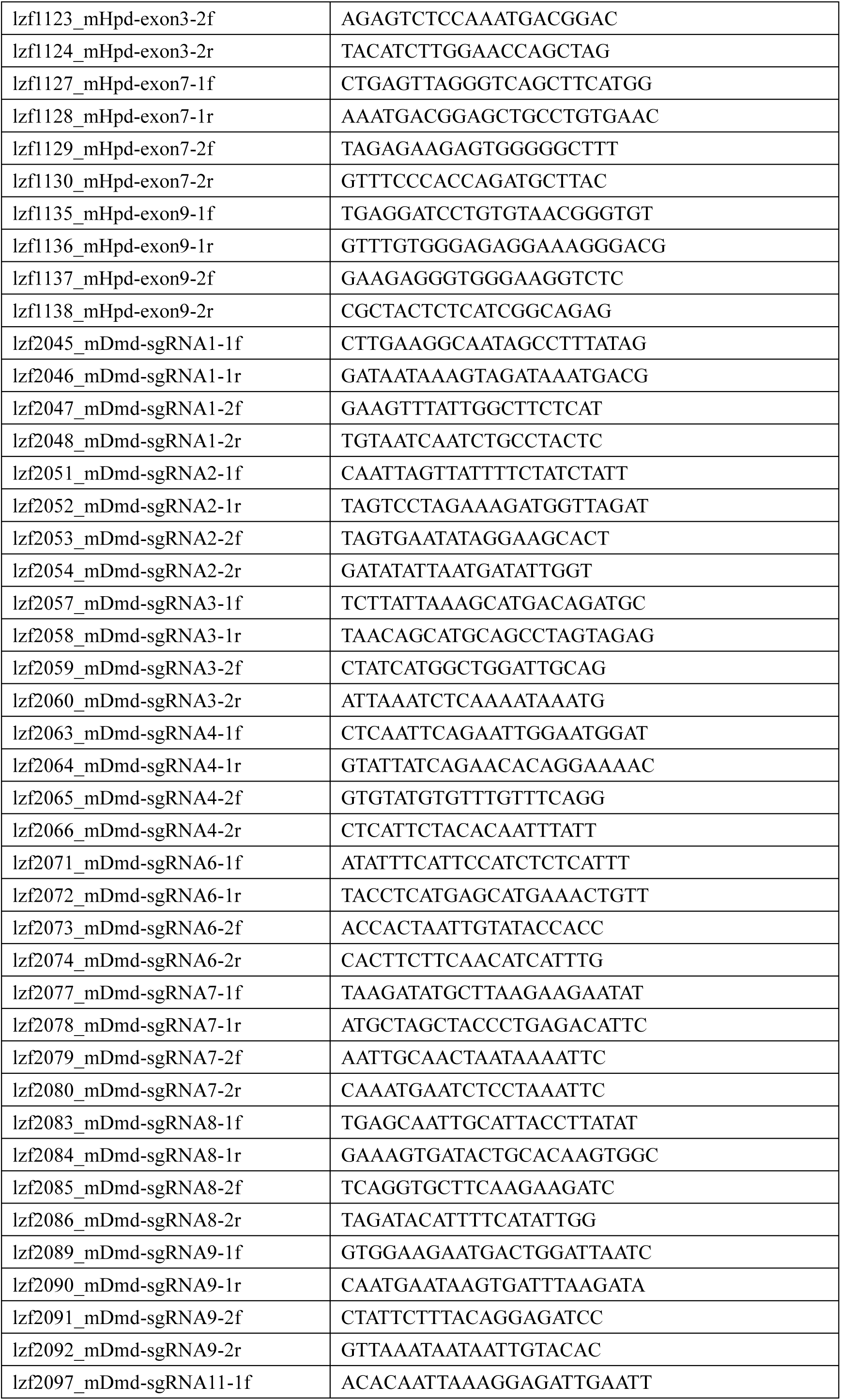
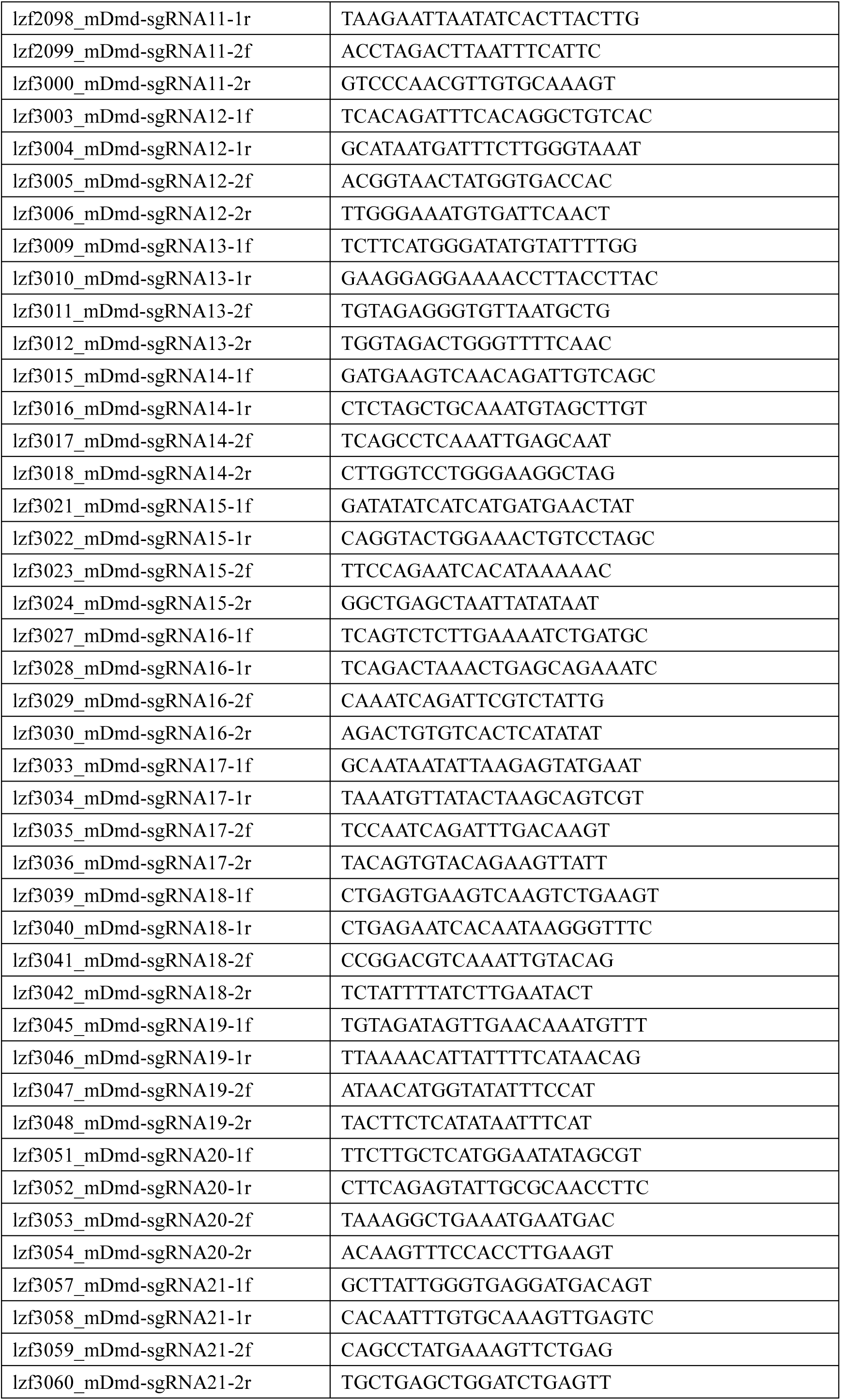
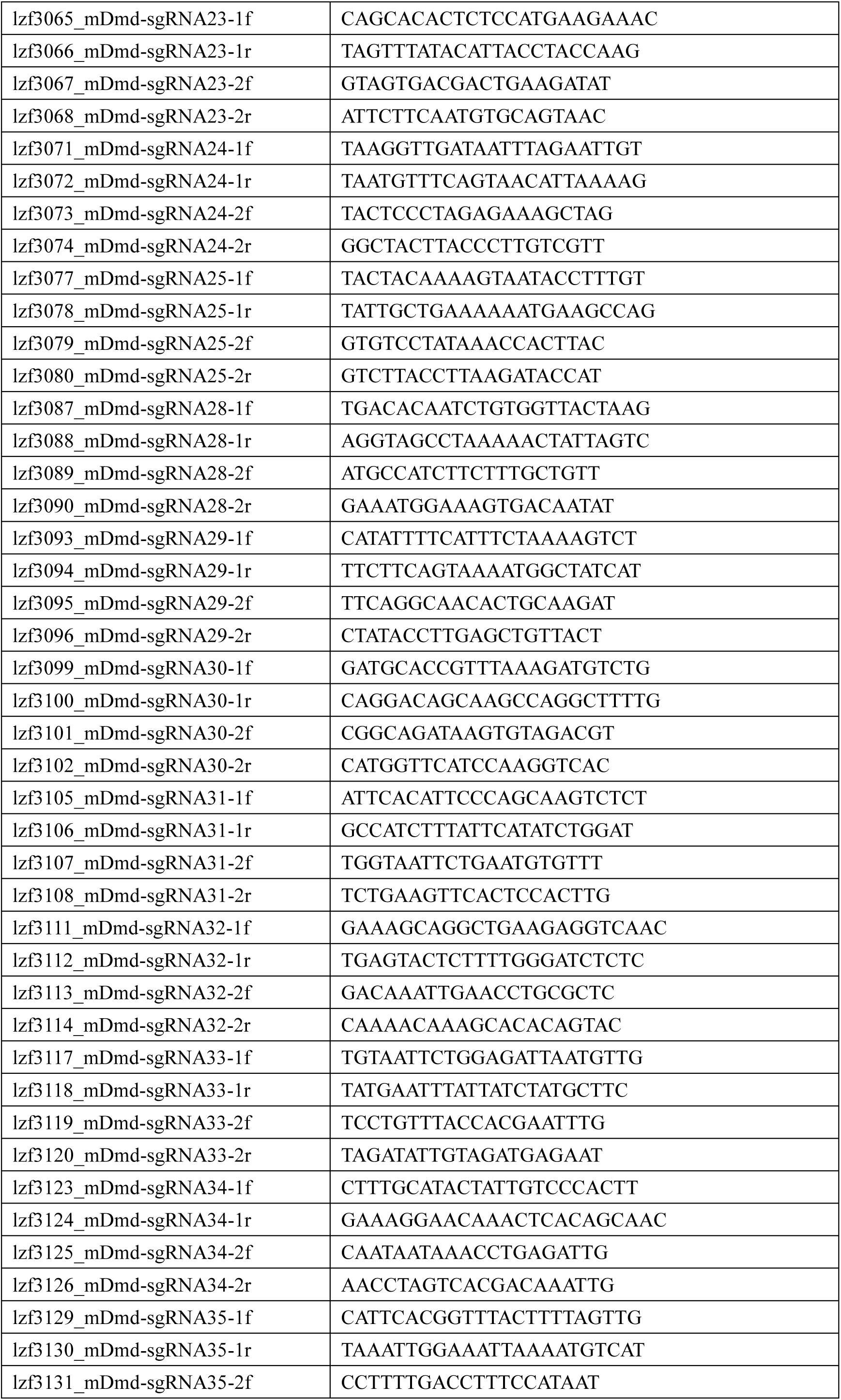
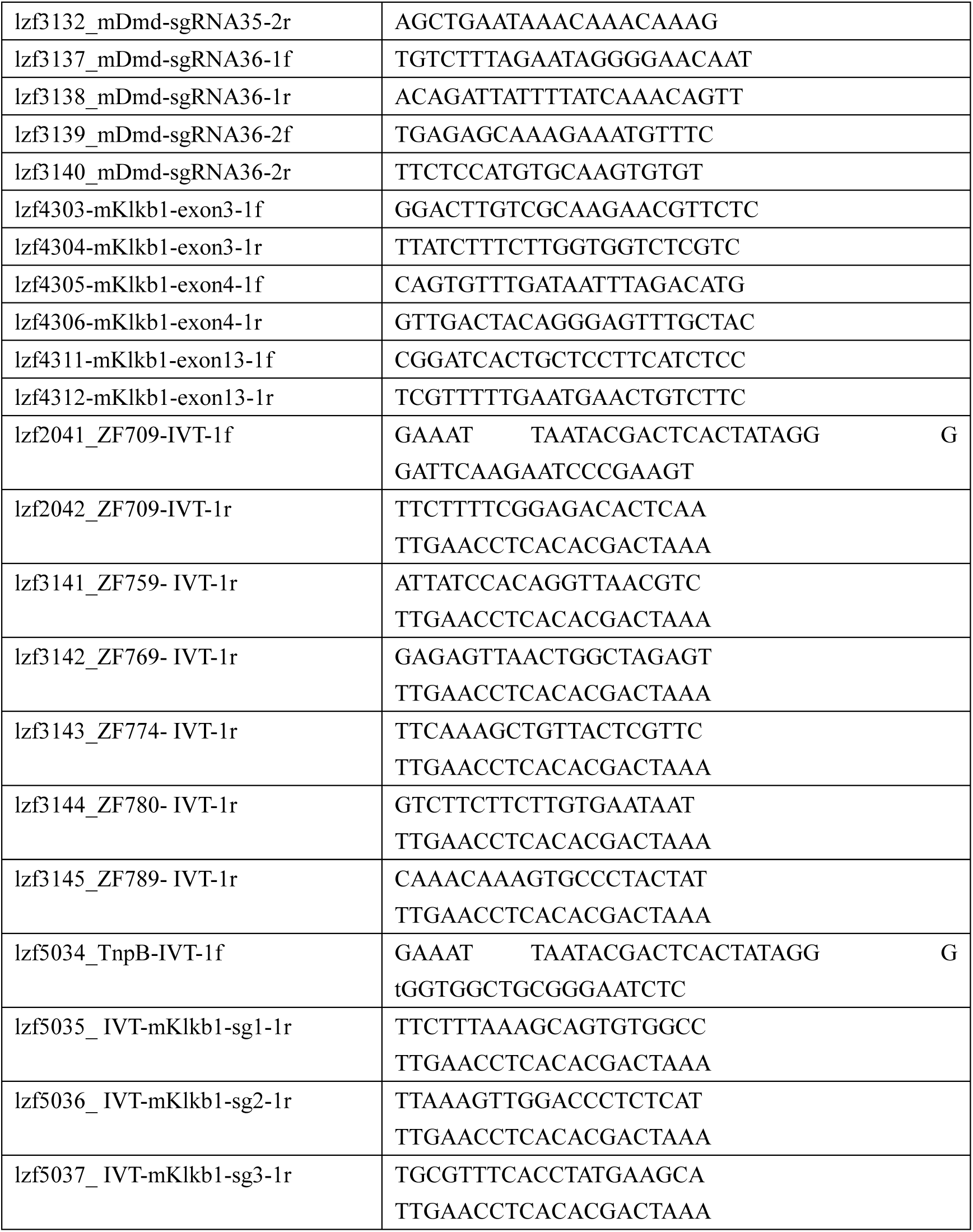
PCR and IVT primers used in this study.

**Supplementary Table S3.**
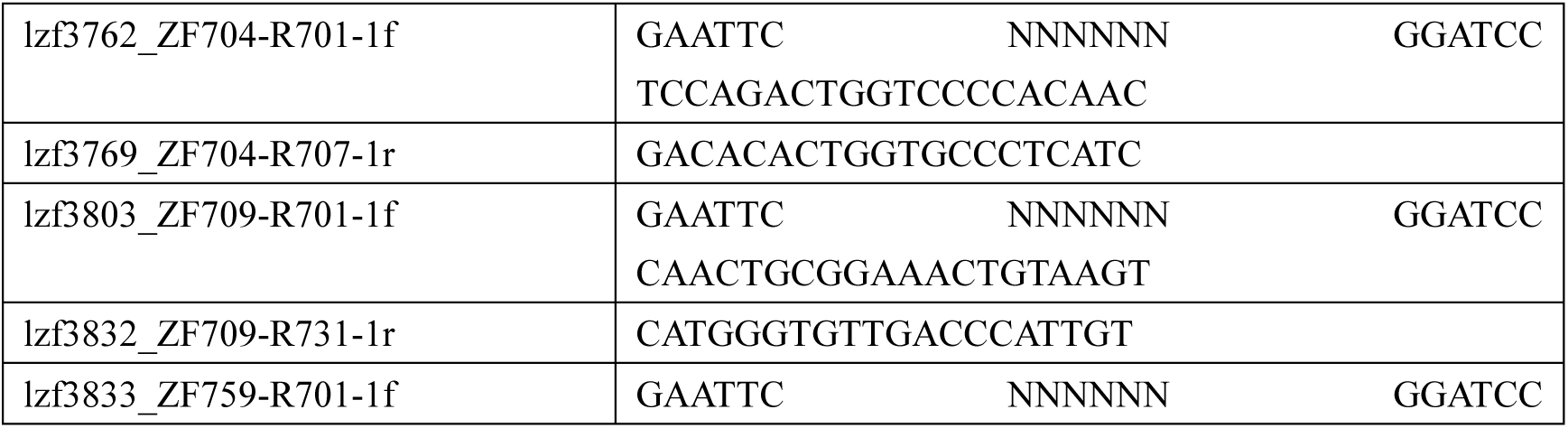
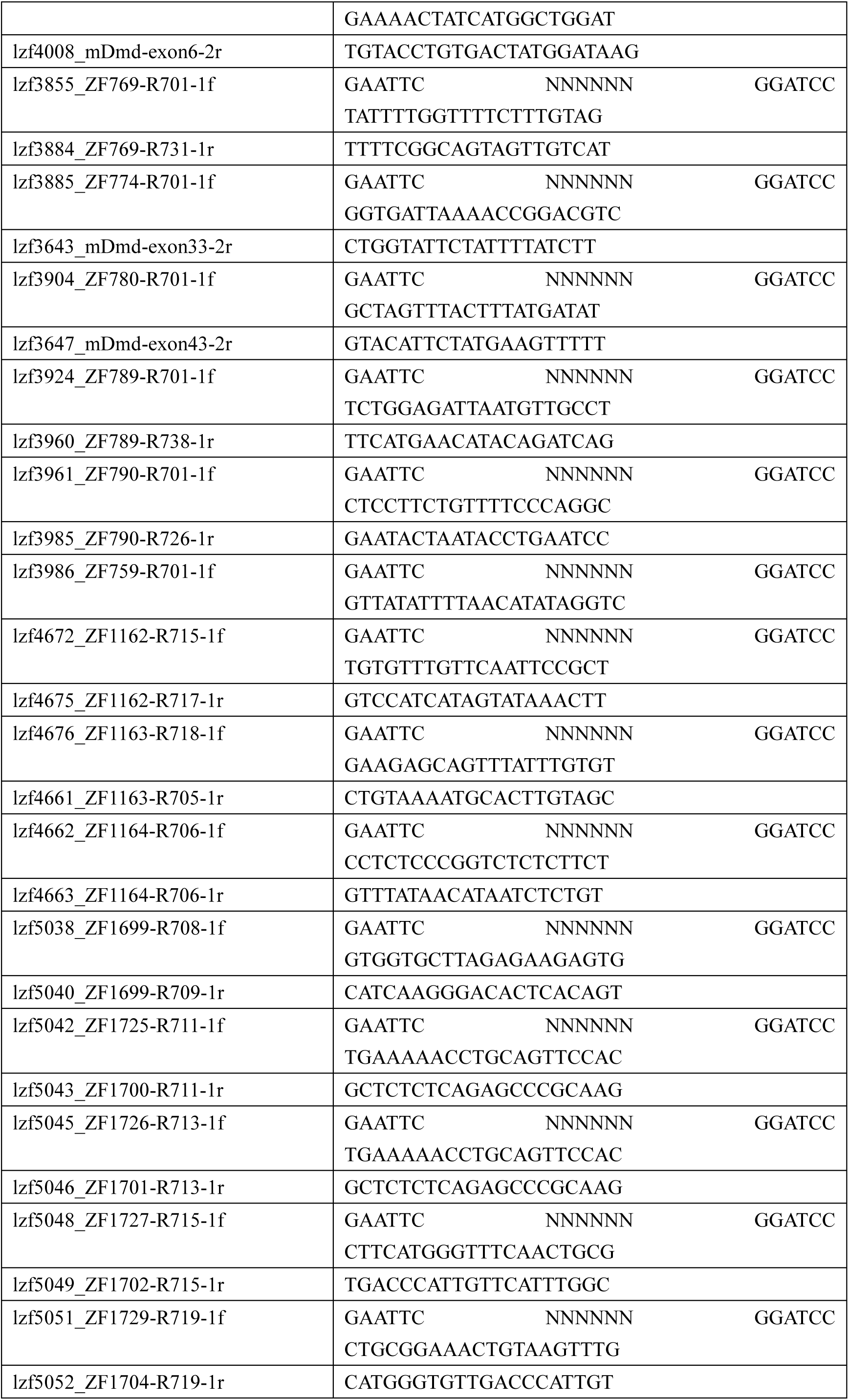
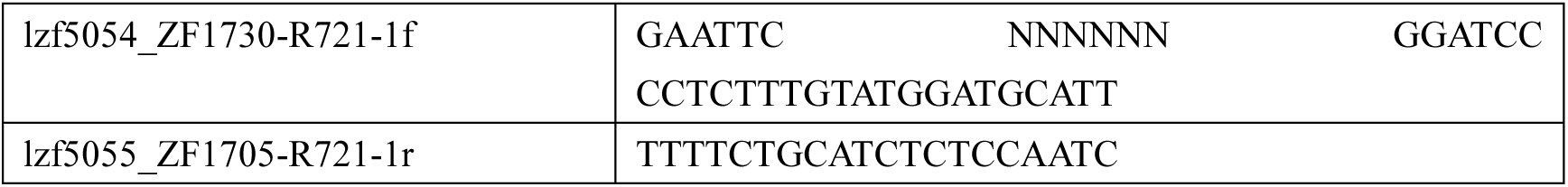
NGS primers used in this study.

