## Supplementary figures and images for "Engineering transposon-associated TnpB-ωRNA system for efficient gene editing and disease treatment in mouse"

### Supplementary figures

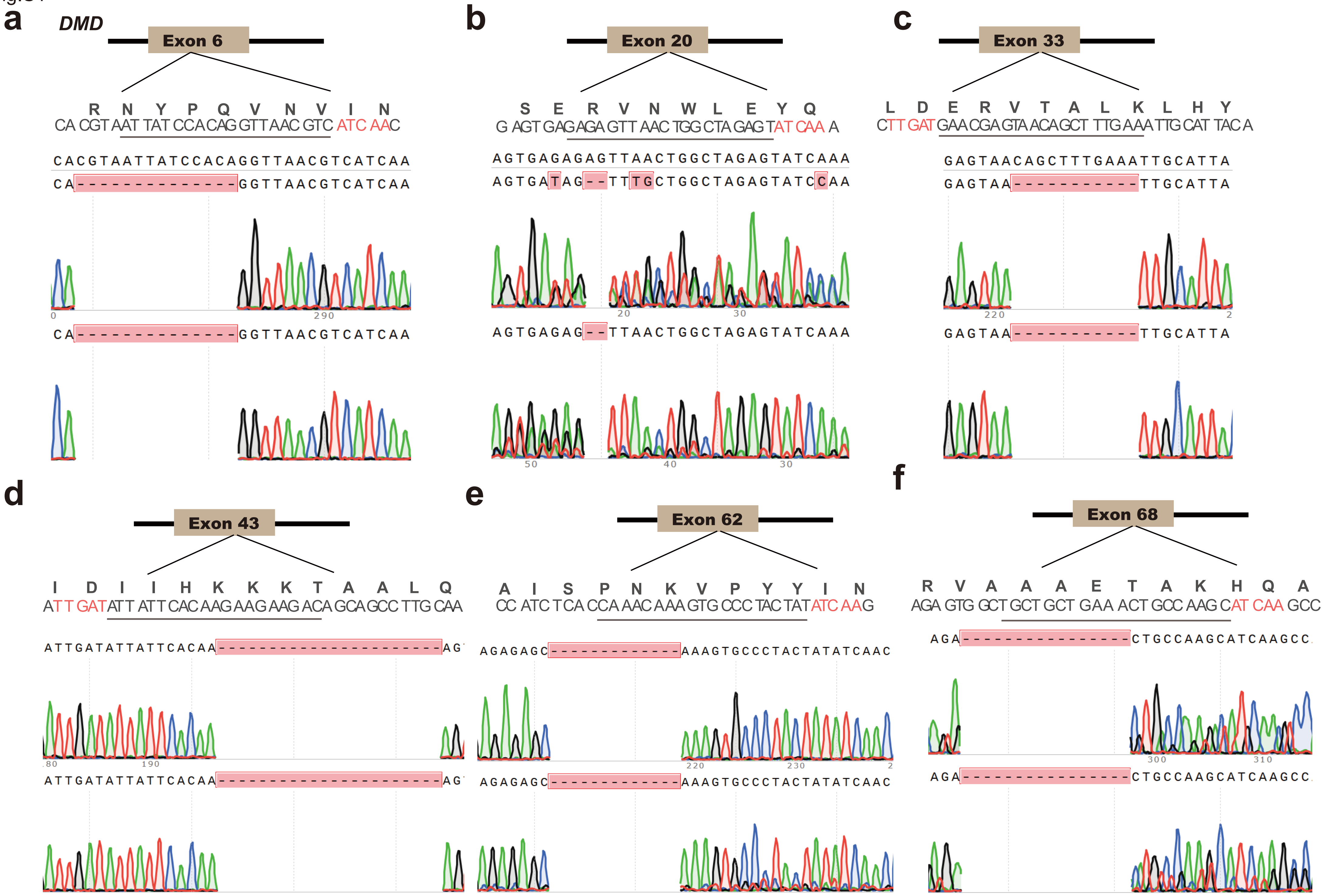

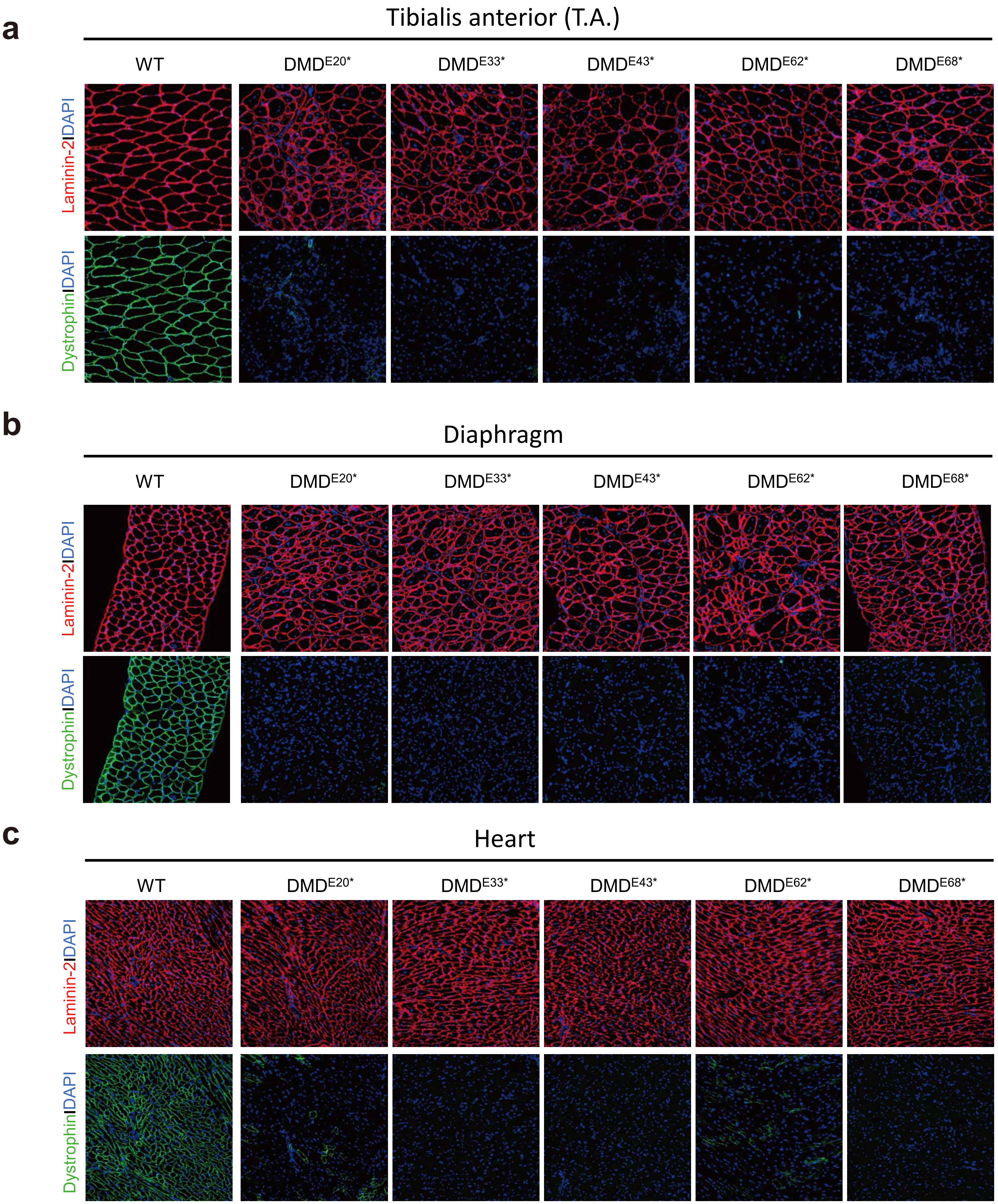

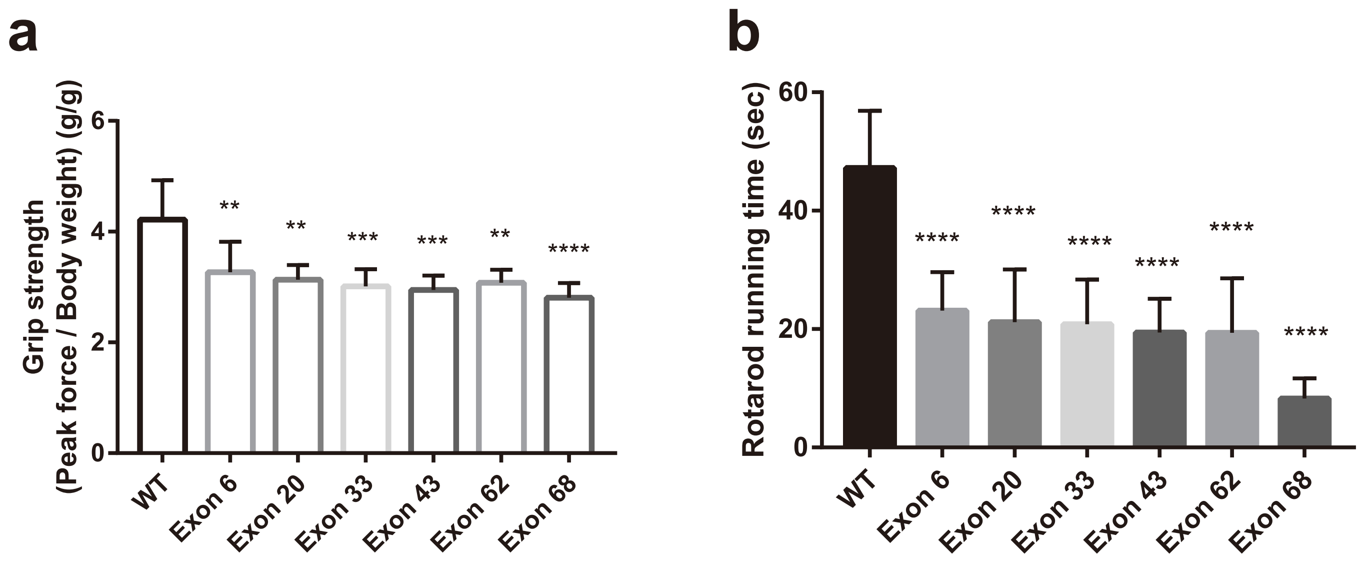

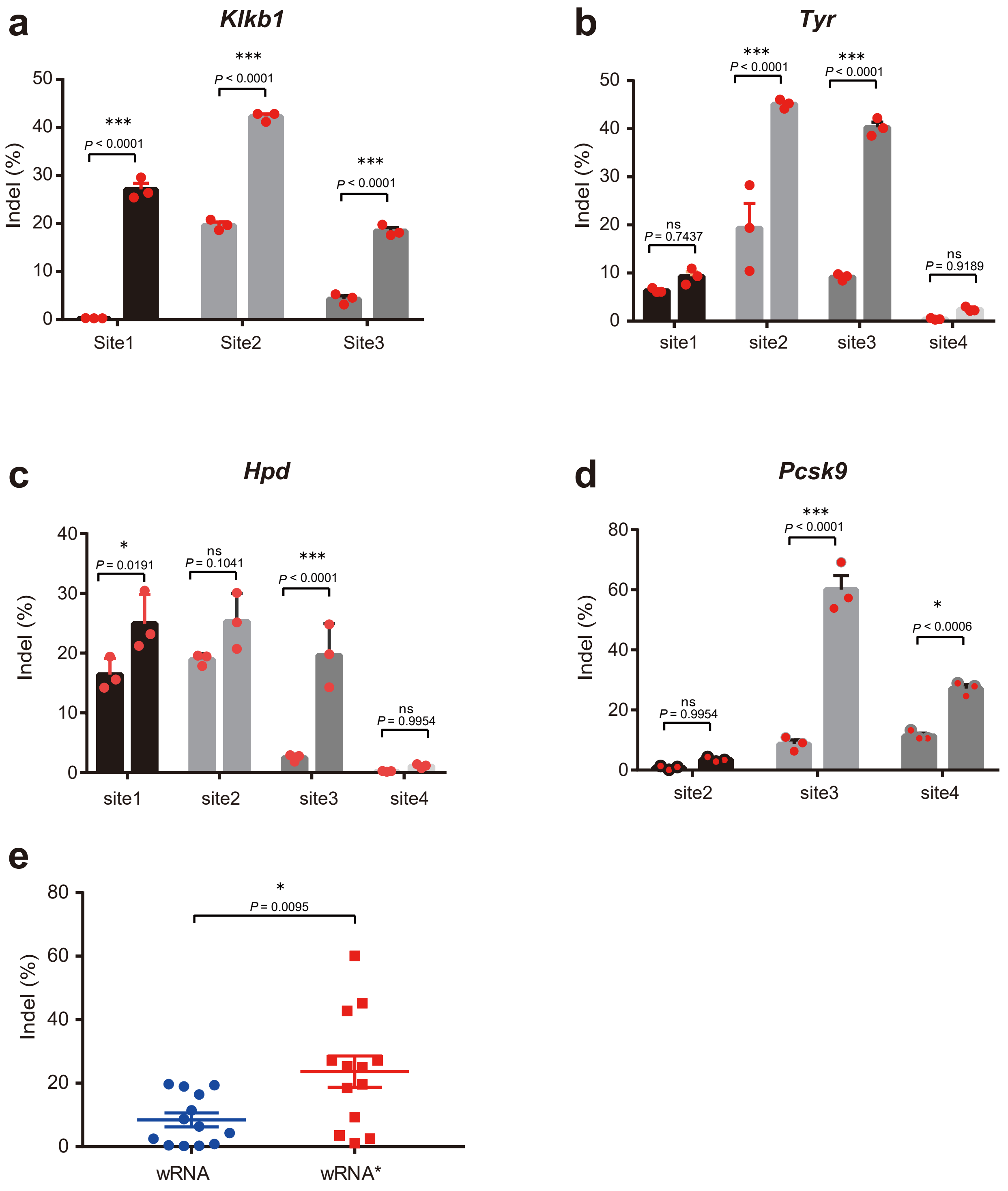

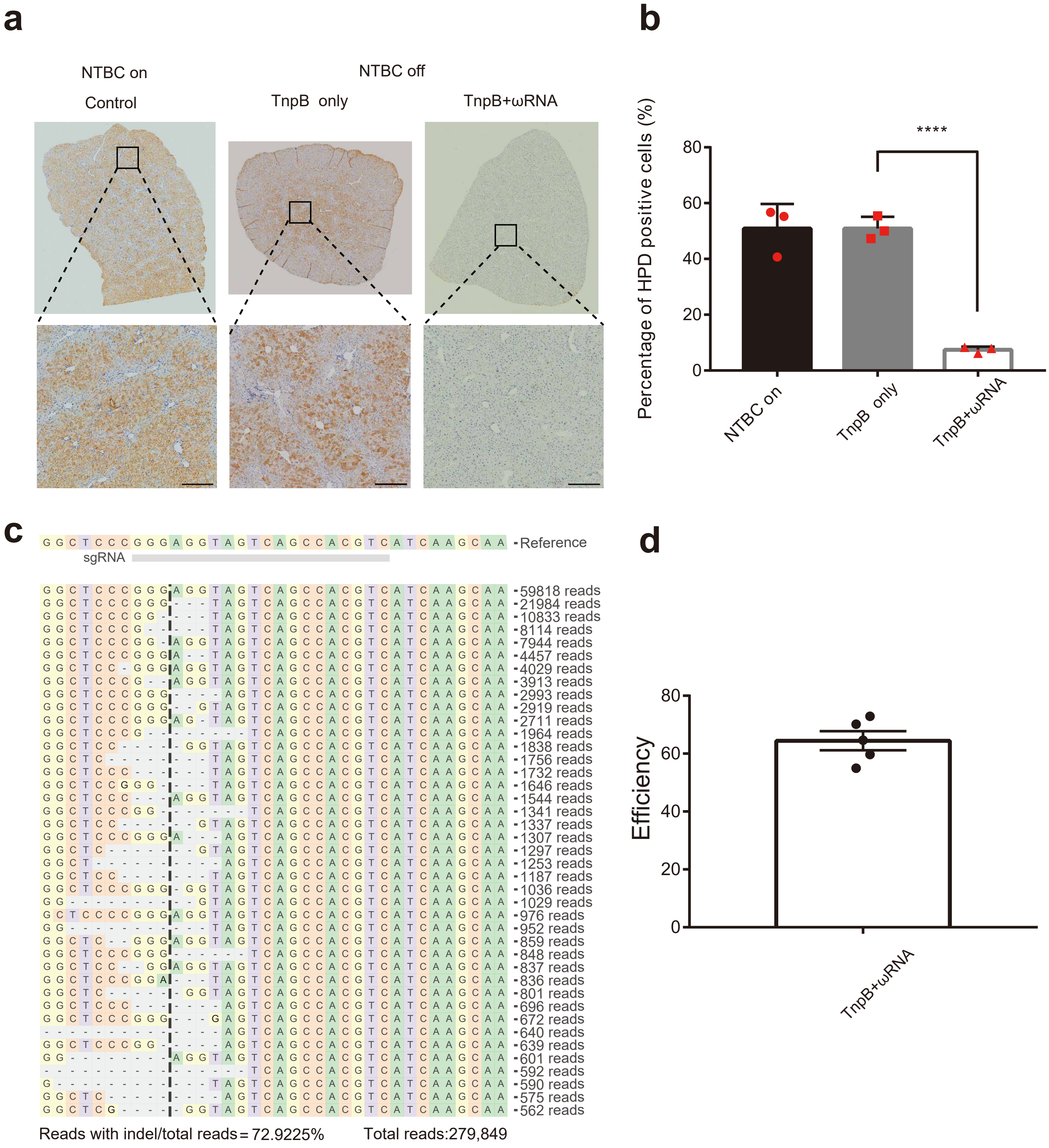

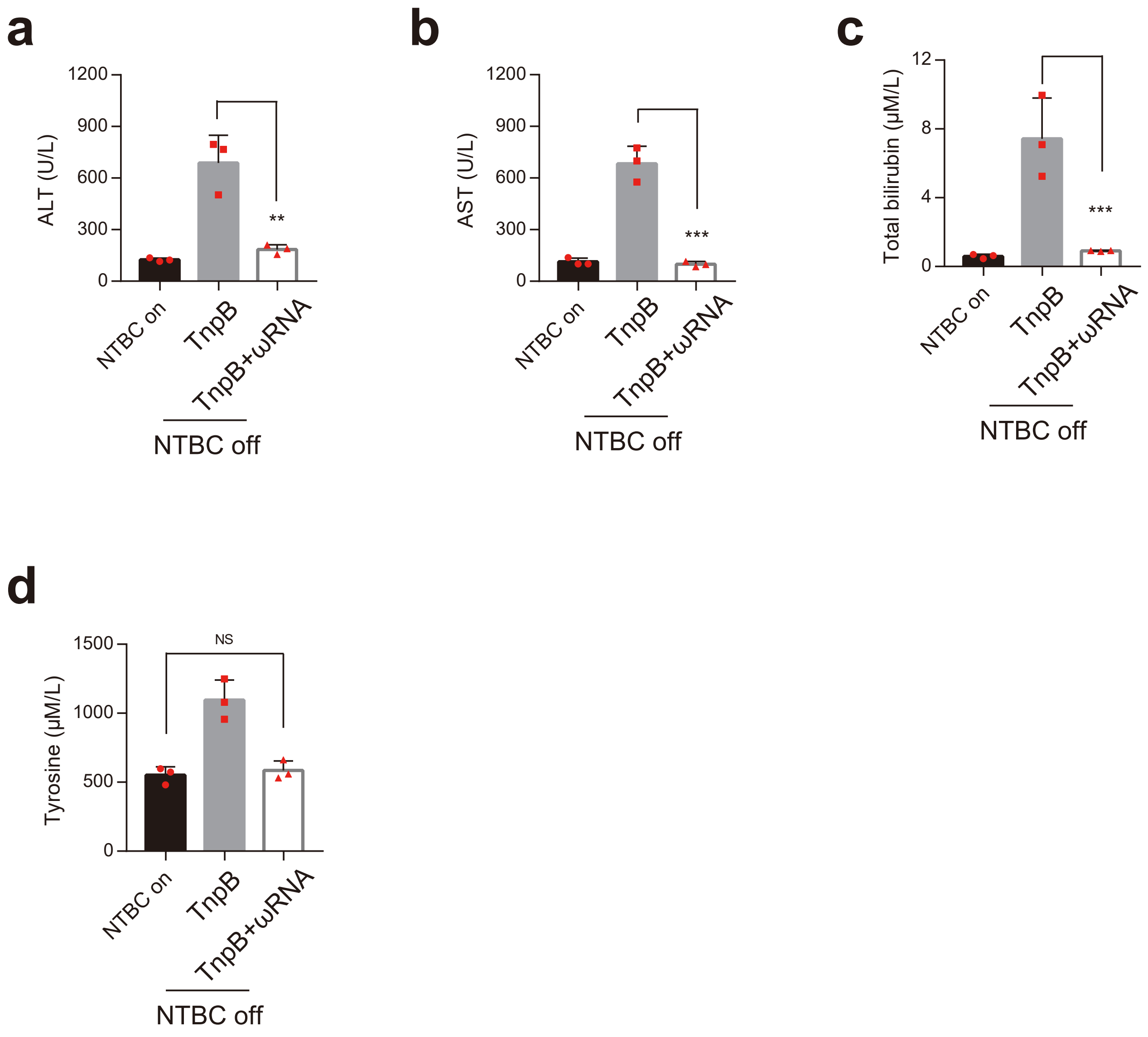
